# Integration of physiologically relevant photosynthetic energy flows into whole genome models of light-driven metabolism

**DOI:** 10.1101/2021.11.04.467239

**Authors:** Jared T. Broddrick, Maxwell A. Ware, Denis Jallet, Bernhard O. Palsson, Graham Peers

## Abstract

Characterizing photosynthetic productivity is necessary to understand the ecological contributions and biotechnology potential of plants, algae, and cyanobacteria. Light capture efficiency and photophysiology have long been characterized by measurements of chlorophyll fluorescence dynamics. However, these investigations typically do not consider the metabolic network downstream of light harvesting. In contrast, genome-scale metabolic models capture species-specific metabolic capabilities but have yet to incorporate the rapid regulation of the light harvesting apparatus. Here we combine chlorophyll fluorescence parameters defining photosynthetic and non-photosynthetic yield of absorbed light energy with a metabolic model of the pennate diatom *Phaeodactylum tricornutum*. This integration increases the model predictive accuracy regarding growth rate, intracellular oxygen production and consumption, and metabolic pathway usage. Additionally, our simulations recapitulate the link between mitochondrial dissipation of photosynthetically-derived electrons and the redox state of the photosynthetic electron transport chain. We use this framework to assess engineering strategies for rerouting cellular resources toward bioproducts. Overall, we present a methodology for incorporating a common, informative data type into computational models of light-driven metabolism for characterization, monitoring and engineering of photosynthetic organisms.

## Introduction

There is great interest in characterizing light-driven metabolism due to the ecological importance and engineering potential of phototrophic microorganisms and plants. Oxygenic photosynthesis utilizes light energy to generate an oxidized protein complex capable of extracting electrons from water at Photosystem II, while concurrently re-energizing the extracted electron to reduce NADP^+^ at Photosystem I. These “light harvesting” reactions drive electron transport, ATP generation, and subsequent CO_2_ fixation through the Calvin-Benson-Bassham cycle in addition to the other energy consuming reactions throughout the cell.

Light absorption by a photosynthetic cell is not constant. Light fluxes can vary across the day and due to local ecological or climatological features. It is common for photosynthetic microorganisms to absorb more photons than what can be utilized by metabolism during these natural fluctuations in sunlight. If this energy is not dissipated, it results in over reduction of the photosynthetic electron transport chain (ETC). Reactive oxygen species are then formed, causing oxidative damage to proteins, lipids, and nucleic acids (Niyogi, 2000; Dietz *et al*, 2016). When the damage resulting from excess light capture results in a decrease in photosynthetic efficiency it is termed photoinhibition. The photosystem II (PSII) D1 subunit is the primary photoinhibition target in the photosynthetic ETC (Edelman & Mattoo, 2008). A complex repair cycle characterized by removal, degradation, and *de novo* synthesis is constitutively active to counter this damage and it is energetically expensive (Nixon *et al*, 2005). To prevent photoinhibition, excess energy can be dissipated upstream of the photosynthetic ETC complexes via a variety of mechanisms encompassing nonphotochemical quenching (NPQ), which harmlessly converts excitation energy to heat (Nicol *et al*, 2019). While this protects the photosynthetic system from oxidative stress, it also reduces the overall efficiency of light-biomass conversion.

Here we coin the term Excess Electron Transport (EET) as an additional important physiological feature at the intersection of photophysiology and bioengineering. This comprises several components that act either as shunts within the ETC (Jallet *et al*, 2016b; Ware *et al*, 2020) or downstream of the photosynthetic machinery within the broader metabolic network. It relieves over reduction of the photosynthetic ETC by dispelling electrons generated by excess light (Jallet *et al*, 2016b). Usually these reactions are considered metabolically “futile” as the electrons are deposited on elemental oxygen to generate water, for instance. However, they are important in relieving photosynthetic ETC over-reduction. There is interest in the bioengineering field to redirect these electrons away from metabolic futility towards bioproducts of interest, while maintaining the beneficial effects on ETC redox balance (Levering *et al*, 2015; Lassen *et al*, 2014). Harnessing excess reductant can convert endogenous carbon sinks, such as carbohydrates, into more energy dense products such as lipids. Indeed, recently it was shown engineered reductant sinks can actually increase carbon fixation and overall photosynthetic efficiency (Santos-Merino *et al*, 2021). Thus, downregulating evolutionarily beneficial processes for photosynthetic individuals in favor of mass culture productivities offers promising avenues for increasing bioproduct and biofuel efficiency. Quantitative characterization of the push-pull of light capture upstream and dissipation in the metabolic network downstream of the photosynthetic ETC would enable design and optimization of these engineered reductant sinks.

Properly accounting for EET facilitates this bioprocess optimization and provides insight into photoprotection strategies. Previous work in photosynthetic microorganisms (diatoms and green algae) used photophysiology parameters derived from chlorophyll fluorescence measurements to estimate EET (Wagner *et al*, 2006). In this previous framework, EET was calculated as the difference between the total absorbed photons and the excitation energy required for biomass production and cellular maintenance. Additionally, the fraction of total absorbed light energy lost upstream of the photosynthetic ETC was estimated using chlorophyll fluorescence data, which have long been employed to assess phototrophic physiology (Krause & Weis, 1991).

Chlorophyll fluorescence primarily quantifies the fate of absorbed light energy directed to PSII; however, there is evidence of contributions from photosystem I (PSI) as well (Giovagnetti *et al*, 2015; Pfündel *et al*, 2013). This excitation energy has three primary fates: it can perform photochemistry at PSII; it can be dissipated as heat through NPQ processes; or it can be dissipated by other, less well characterized non-radiative and fluorescence processes (NO). All of these can be quantified through the use of pulse amplitude modulation (PAM) chlorophyll fluorimetry (Kramer *et al*, 2004). When these values are normalized to the total excitation energy routed to PSII, they are annotated as the quantum yields Y(II), Y(NPQ) and Y(NO), respectively, the sum of which is always one. These techniques have unveiled the diverse photoprotective strategies employed by photosynthetic microorganisms to include extensive NPQ in the diatom *Phaeodactylum tricornutum* (Lavaud *et al*, 2002). However, these important aspects of photosynthesis have not been integrated in to models of total cellular metabolism.

Constraint-based modeling coupled with flux balance analysis (FBA) has successfully been employed to characterize and engineer a wide range of biological systems (O’Brien *et al*, 2013). Constraint-based modeling relies on a reconstruction of the metabolic content of the organism of interest. The resulting computational framework, known as a genome scale model (GEM), can then be used to compute a variety of cellular phenotypes. There have been several, recent advances in the metabolic modeling of photosynthetic organisms to include cyanobacteria (Broddrick *et al*, 2016, 2019b), green algae (Zuñiga *et al*, 2017; Chang *et al*, 2011), and diatoms (Levering *et al*, 2016). Recent modeling in the diatom *Phaeodactlyum tricornutum* quantified growth rates, excitation energy partitioning between the photosystems and cross-compartment energetic coupling of the chloroplast and mitochondrion (Broddrick *et al*, 2019a). However, that study used simplified assumptions regarding light harvesting, possibly affecting the accuracy of absolute fluxes predicted by the model. The metabolic network that underpins GEMs is assembled from the reactant and product stoichiometry of biochemical reactions; thus, it should be feasible to couple the representation of chlorophyll fluorescence parameters as a fraction of light energy routed to PSII as a stoichiometrically balanced biochemical equation. Such a framework would enable the explicit integration of chlorophyll fluorescence data and EET as a constraint on photosynthetic metabolic processes towards an increased understanding of photoacclimation, photoprotection and bioengineering of phototrophic metabolism.

## Results

### Cell physiology of P. tricornutum at low and high light

*P. tricornutum* was acclimated and cultured at a high light irradiance of 600 µmol photons m^−2^ s^−1^ (HL, n=4) and a low light irradiance of 60 µmol photons m^−2^ s^−1^ (LL, n=3). The range of growth rates for *P. tricornutum* was 0.026-0.029 (n=3) and 0.052-0.053 (n=4) hr^−1^ at LL and HL, respectively. Cell volumes for cultures grown at both light levels differed by approximately 10% (202 ± 43 versus 184 ± 47 µm^3^ at HL (n=94) and LL (n=46), respectively). Dry cell weight was also similar between the cultures (Table 1).

**Table 1.** Physiology parameters of *P. tricornutum* acclimated to low and high light.

| | Growth rate<br>($\text{h}^{-1}$) | Cell volume<br>($\mu\text{m}^3$) | Cell weight<br>(pgDW/cell) | Chla<br>(pg/cell) | Chlc<br>(pg/cell) | Chla/Chlc | Total chl<br>(LL:HL) |
| --- | --- | --- | --- | --- | --- | --- | --- |
| Low Light | 0.026-0.029 | $184 \pm 47$ | $19.1 \pm 2$ | $0.384 \pm 0.006$ | $0.073 \pm 0.001$ | 6.0 | 2.8 |
| High Light | 0.052-0.053 | $202 \pm 43$ | $20.4 \pm 1.4$ | $0.142 \pm 0.014$ | $0.024 \pm 0.003$ | 5.2 | |

There were differences in the chlorophyll content of cells adapted to different light regimes, as is typical for microalgae (Falkowski & Owens, 1980). Total chlorophyll (chlorophyll *a* (chla) and chlorophyll *c* (chlc)) at LL was 2.8-fold higher than at HL (Table 1) and resulted in a 3-fold increase in the cell normalized absorption coefficient (a*_cell_, Fig. 1A, B); suggesting light capture efficiency remained constant at both acclimated light conditions. The chla to chlc ratio varied slightly from 6.0 at LL to 5.2 at HL. Overall, the chlorophyll content per cell was consistent with previous observations of photoacclimation in *P. tricornutum* (Nymark *et al*, 2009; Broddrick *et al*, 2019a).

**Figure 1.**
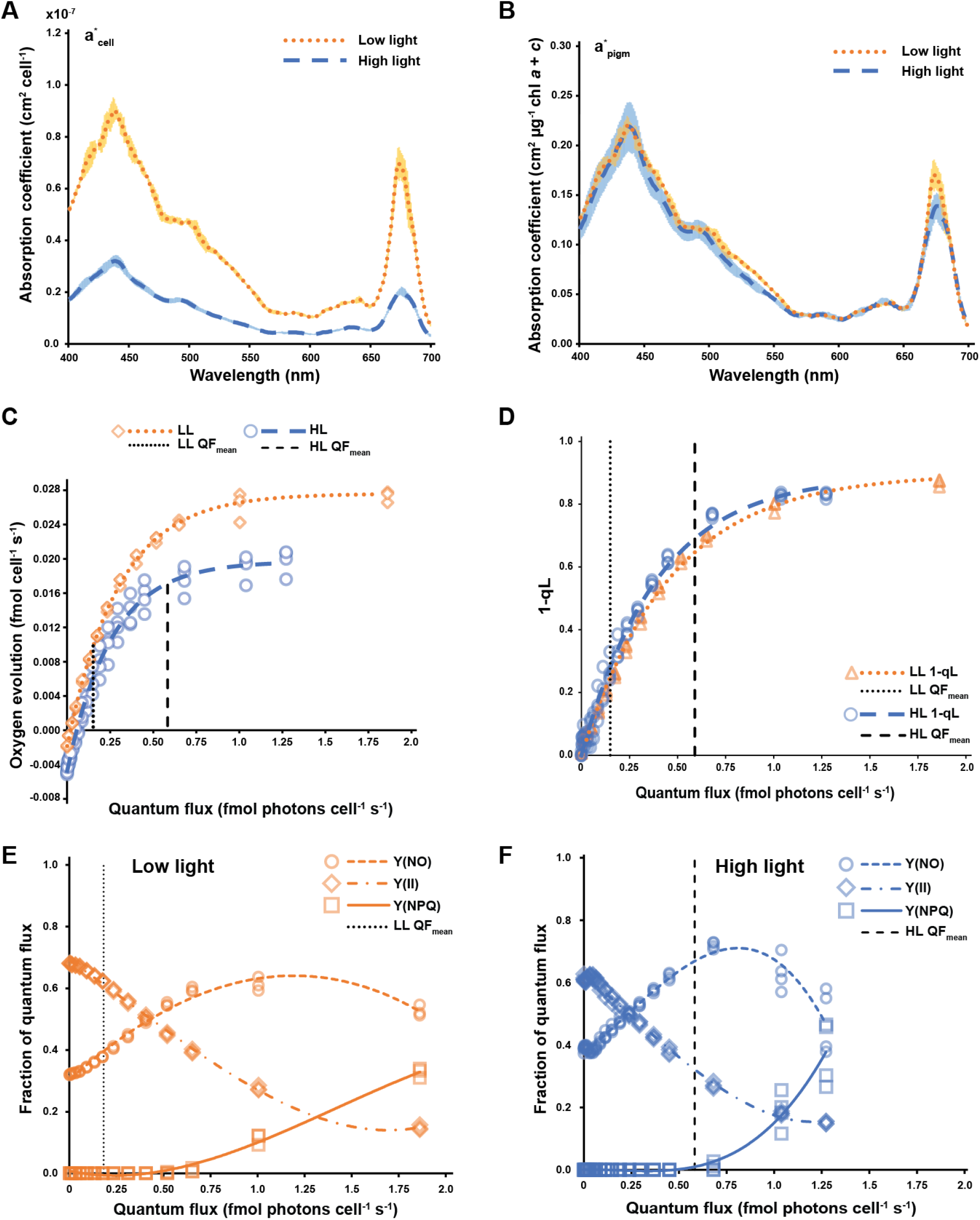
Photophysiology of *P. tricornutum* acclimated to low and high light. (A) Cell-specific absorption coefficient (B) Pigment-specific absorption coefficient. The pigment mass includes chlorophyll *a* and chlorophyll *c*. Shaded areas represent one standard deviation from the mean (HL: n=4, LL: n=3). (C) Cell-specific PO versus QF curve. (D) Fraction of closed reaction centers (1-qL) versus QF curve. (E) Chlorophyll fluorescence parameters vs. quantum flux for cells acclimated to low light. (D) Chlorophyll fluorescence parameters vs. quantum flux for cells acclimated to high light. Vertical dashed lines represent the mean quantum flux received by the cultures at the experimental irradiance. Abbreviations and definitions: LL: low light, HL: high light, QF: quantum flux, Y(II): quantum efficiency of photosystem II, NPQ: non-photochemical quenching, Y(NO): unregulated, non-radiative dissipation of excitation energy. Data based on n=3 biological replicates for LL and n=4 biological replicates for HL.

### Photophysiology of P. tricornutum at low and high light

Using a rapid light curve (RLC) protocol (Jallet *et al*, 2016a) we concurrently determined chlorophyll fluorescence parameters and oxygen evolution. The genome-scale model (GEM) was constrained with the photon uptake rate (quantum flux, QF) and the oxygen evolution rate in a manner similar to recent modeling efforts in cyanobacteria and diatoms (Broddrick *et al*, 2019a, 2019b). Accounting for the change in absorbed QF across the culture due to cellular self-shading, we report both the maximum QF (QF_max_) and the mean QF (QF_mean_), representative of the highest and the average photon capture rate across the full path length. As PAM measurements require high cell densities to generate sufficient fluorescence signal, we used a similar approach to account for cell shading in the PAM sample cuvette (Broddrick *et al*, 2019b). This calculated QF_mean_ was used as the independent variable for P_O_ vs. QF curves as well as plots of chlorophyll fluorescence parameters vs. QF.

The growth rate differences between HL and LL acclimated cultures were largely attributed to photophysiology differences between the two conditions. Despite the HL acclimated cells absorbing 3.3 times more photons than the LL acclimated cells, the HL maximum oxygen evolution rate was only 1.3-fold higher (Po_max_, Table 2). However, this ratio increased to 1.7-fold at the mean oxygen evolution rate (Table 2, Po_mean_), quantifying the impact of self-shading from the increased pigment content at LL on overall productivity. The P_O_ vs. QF curve initial slopes were 9.5×10^−2^ and 9.8×10^−2^ mol O_2_ mol photon^−1^ for HL and LL, respectively. This similarity in values was also observed with the initial slope of the total chlorophyll (chla+chlc) normalized-P_O_ vs. PAR curves which was determined to be 2.6×10^−4^ mol O_2_ mol photon^−1^ m^2^ mgChl^−1^ for both HL and LL (Fig. S1).

**Table 2.** Comparison of photophysiology in *P. tricornutum* acclimated to low and high light.

|  | QF<sub>max</sub><sup>*</sup> | QF<sub>mean</sub><sup>*</sup> | P<sub>O</sub><sub>max</sub><sup>†</sup> | P<sub>O</sub><sub>mean</sub><sup>†</sup> | Fv/Fm | Y(II)<sup>‡</sup> | Y(NPQ)<sup>‡</sup> | 1-qL<sup>‡</sup> | P<sub>O</sub><sub>max</sub><br>(HL:LL) | P<sub>O</sub><sub>mean</sub><br>(HL:LL) |
| --- | --- | --- | --- | --- | --- | --- | --- | --- | --- | --- |
| Low Light | 0.23 | 0.15 | 0.014± 0.000 | 0.010 ± 0.000 | 0.68 | 0.63 | 0.00 | 0.25 | 1.3 | 1.7 |
| High Light | 0.77 | 0.59 | 0.018± 0.002 | 0.017± 0.002 | 0.63 | 0.32 | 0.00 | 0.69 |  |  |

<sup>\*</sup> fmol photons cell<sup>-1</sup> s<sup>-1</sup>
<sup>†</sup> fmol O<sub>2</sub> cell<sup>-1</sup> s<sup>-1</sup>
<sup>‡</sup> Value at QF<sub>mean</sub>

Overall, trends in the chlorophyll fluorescence parameters plotted against quantum flux (PAM vs. QF) were similar between the two light regimes (Fig. 1D-F). The effective quantum yield of PSII, [Y(II)], decreased rapidly with increasing QF for both light regimes. Non-photochemical quenching, contributed almost no excitation energy dissipation at either HL or LL (Table 2 and Fig. 1E, F). As Y(II) accounts for the fraction of QF performing photochemistry and Y(NPQ) is the fraction of QF lost as heat, the balance, Y(NO), accounts for the remaining fraction of QF that is dissipated in unregulated, non-radiative processes. Y(NO) accounted for approximately 68% and 37% of PSII-directed excitation energy at QF_mean_ for HL and LL, respectively. The fraction of closed reaction centers (1-qL) is a proxy for the redox state of the plastoquinone pool and it was almost identical across the entire QF range for HL and LL acclimated samples (Fig. 1D). Additionally, NPQ had similar activation profiles in both LL and HL acclimated cultures. For both LL and HL acclimated cultures, Y(NPQ) activated at a QF of approximately 0.7 fmol photons cell^−1^ s^−1^, 1-qL values of 0.70-0.75 and approximately 90% of their respective maximum photosynthetic rates (Fig. S2).

The cell densities used, and photosynthetic rates observed, at high light suggested carbon limitation in the samples during analysis. Initially, we did not supplement the PAM samples with bicarbonate as we were trying to assess the photophysiology of the experimental culture, which was only sparged with air. However, to assess the possibility of carbon limitation and its influence on light capture efficiencies, we repeated the P_O_ vs. QF and PAM experiments supplementing the samples with 5 mM bicarbonate. The resulting differences in P_O_ vs. QF for *P. tricornutum* suggested a 15% underestimation of oxygen evolution capacity at QF_mean_ for the HL acclimated condition, but additional bicarbonate supplementation had no impact on the LL condition. (Fig. S3, B). Additionally, PAM results were comparable between samples with and without added bicarbonate (Fig. S3C), suggesting the carbon limitation was minor enough to not affect quantum efficiency.

### Simulating photoautotrophic growth of P. tricornutum at low and high light

We simulated photoautotrophic growth at low and high light by translating the photophysiology results into modeling constraints (Broddrick *et al*, 2019a, 2019b). The photon uptake rate for the model was derived from a*_cell_ coupled with the experimental PAR intensity and the emission spectrum of the fluorescent light used during culturing. This value, equivalent to QF, was used to determine the oxygen evolution rate of the culture (using the curves in Fig. 2), which constrained the oxygen exchange reaction for the simulations.

**Figure 3.**
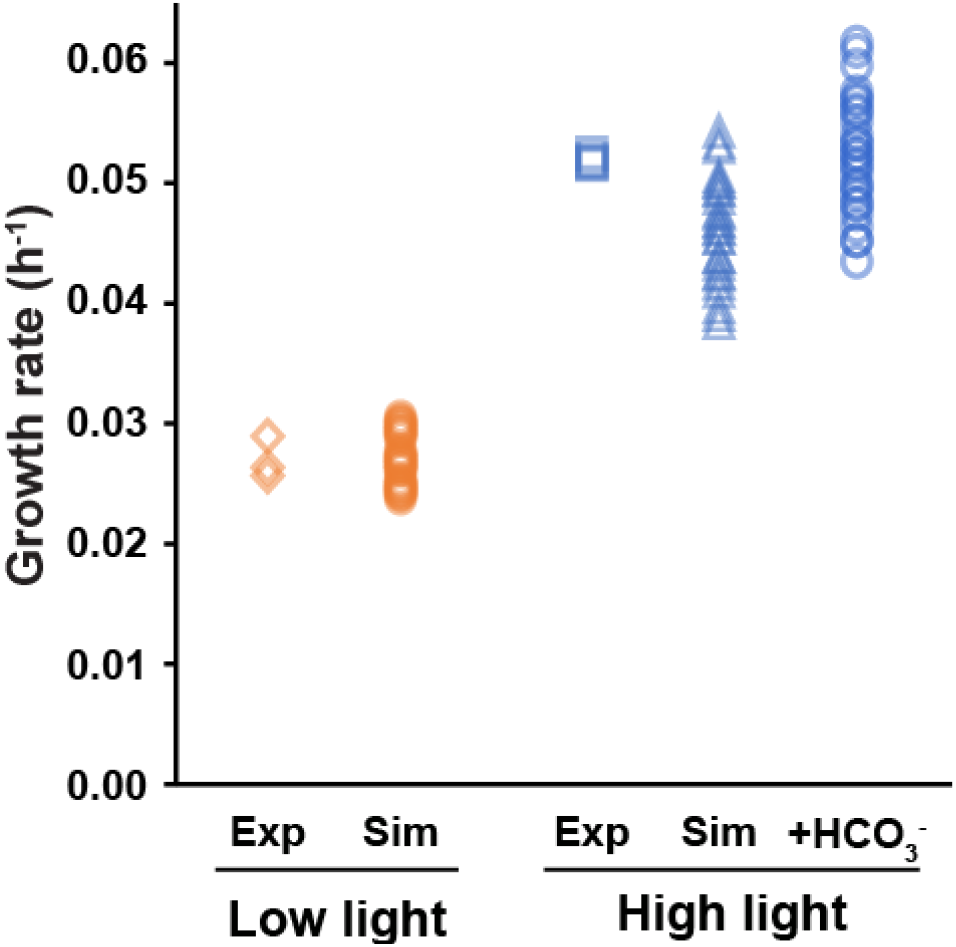
Experimental versus simulated growth rates for *P. tricornutum* acclimated to low and high light. Experimental values represent the growth rate for independent biological replicates (HL: n=4, LL: n=3). For the simulation values, the data points represent the simulated growth rate using the mean, +1 standard deviation and −1 standard deviation of the O_2_ vs QF curves, a^*^_cell_ and cell dry weight (n=27 parameter combinations). Abbreviations. Exp: experimental, Sim: simulated, +HCO_3_^−^: simulated, bicarbonate spiked.

Next, we incorporated the chlorophyll fluorescence data with our modeling construct. GEMs are built using biochemical reaction stoichiometry. Chlorophyll fluorescence parameters are normalized as fractional values of excitation energy routed to photosystem II, which can be formulated in a similar manner to stoichiometry in canonical biochemical reactions. Thus, we added a pseudo-reaction to the model that imposes this fractionation between excitation energy lost in the pigment bed (1-F_v_/F_m_), photochemical yield [Y(II)], regulated non-photochemical quenching [Y(NPQ)], and unregulated non-photochemical quenching [Y(NO)] (Fig. S4). When the model simulates photoautotrophic growth, it predicts the excitation energy split between the photosystems to satisfy the reductant and ATP needs for biomass production. The chlorophyll fluorescence parameters apply a constraint on the excitation energy split as only the Y(II) fraction can perform photochemistry at PSII.

A final constraint added to the model accounted for photodamage of the PSII D1 subunit. We determined the D1 damage rate at the experimental irradiance for both LL and HL acclimated cells by comparing the maximum quantum yield of PSII (F_v_/F_m_) with and without lincomycin, a plastid protein synthesis inhibitor (Fig. S5A, B). We determined the D1 damage first-order rate constant to be between 5×10^−4^ and 7×10^−4^ (n=3) and 2.22×10^−2^ and 2.52×10^−2^ (n=3) for LL and HL acclimated samples, respectively. Through Western blot analysis we determined the D1 protein to be approximately 1.1% of total protein for both LL and HL acclimated samples. From these values, we calculated the D1 damage constraint to be 7±2 ×10^−6^ and 2.53±0.51 ×10^−4^ mmol D1 gDW^−1^ hr^−1^ for LL and HL acclimated samples, respectively. These values were added as a maintenance energy reaction to the model requiring 360 mmol ATP and 720 mmol GTP to be consumed at the plastid ribosome to polymerize 1 mmol of D1 protein. A summary of these values can be found in Table S1.

With the suite of photophysiology constraints incorporated into the GEM, we simulated photoautotrophic growth for both light conditions. To account for experimental variability on the model predictions, we simulated growth using a parameter space that included the mean and plus or minus one standard deviation of the experimentally determined P_O_ vs. QF curves (Fig. 1C), a*_cell_ (Fig 1A) and dry cell weight (Table 1). Simulated growth rates were consistent with experimental values (Fig. 2). The model predicted a LL mean growth rate of 0.027 ± 0.002 h^−1^ (n=27 parameter combinations), in good agreement with the experimental range of 0.026-0.029 h^−1^ (n=3). For the HL condition the model predicted a mean growth rate of 0.046 ± 0.004 h^−1^ (n=27 parameter combinations) compared to an experimental range of 0.052-0.053 h^−1^ (n=4). The predicted mean growth rate for the HL acclimated condition was underestimated by approximately 12%; however, adjusting the P_O_ vs. QF curve based on the bicarbonate spiked data (Fig. S3A) resulted in a simulated mean growth rate of 0.052 ± 0.005 (Fig. 6).

### Hierarchy of photophysiology constraints for simulating photoautotrophy

We assessed the relative contribution of various constraints on the accuracy of model predictions, specifically photon uptake, oxygen evolution, photosystem II repair requirements and photochemical yield as constraints on growth rate (Table 3). For the LL simulations, the model was parameterized with the mean values for P_O_ vs. QF, a*_cell_ and dry cell weight. For the HL simulations, we used the mean values for all but P_O_ vs. QF, where we used the bicarbonate spike-adjusted data, as it most accurately represented the photophysiology of the experimental cultures (Fig. 2).

**Table 3:** Hierarchy of constraints for *P. tricornutum* simulations. Abbreviations: *hv* – photon uptake, P_O_ – oxygen evolution rate, D1 – PSII D1 protein repair, Y(II) – PSII quantum yield, DM20% – previous modeling assumption of 20% of absorbed photons lost upstream of the photosystems (Broddrick *et al*, 2019a). Values are given in units of h^−1^.

| *P. tricornutum* |  |  |
| --- | --- | --- |
|  | HL | LL |
| Experimental | 0.052-0.053 | 0.026-0.029 |
| $h\nu$ | 0.136 | 0.038 |
| $P_O$ | 0.052 | 0.027 |
| $h\nu$, $P_O$, D1 | 0.052 | 0.027 |
| $h\nu$, $Y(II)$ | 0.062 | 0.030 |
| $h\nu$, $P_O$, D1, $Y(II)$ | 0.052 | 0.027 |
| $h\nu$, $P_O$, DM20% | 0.052 | 0.027 |

Constraining the photon uptake (*hv*) alone overestimated the growth rate for all conditions (+40% and +160% compared to the experimental mean for LL and HL, respectively – Table 3). This over estimation is to be expected as it assumes all the light captured by the cell is converted into photochemical energy. Using the oxygen evolution rate (P_O_) as the sole constraint resulted in accurate growth rate predictions (within the range of experimentally observed growth rates for both LL and HL). Combining constraints on photon uptake, P_O_ and D1 repair requirements, as well as photon uptake, P_O_, D1 repair, and photochemical yield [Y(II)], did not change the predicted growth rates as compared to P_O_ alone. Constraining the simulations with only the photon uptake and Y(II) resulted in an accurate prediction at LL (+5% compared to the experimental mean). For HL acclimated cells, the prediction was more accurate than photon uptake alone but still over-estimated the growth rate (+18% compared to experimental mean).

### Constraining D1 repair and Y(II) affects metabolic pathway predictions in P. tricornutum

Next, we investigated how the D1 repair requirement and Y(II) constraints affected predictions of metabolic pathway usage. Intracellular reaction fluxes provide insight into the metabolic phenotype and form the basis of GEM-based bioengineering strategies. In previous modeling efforts in *P. tricornutum* acclimated to high light, the predicted rate of intracellular oxygen consumption was higher than that which was observed experimentally (Broddrick *et al*, 2019a). The previous modeling effort assumed up to 20% of the photon flux could be dissipated upstream of the photosystems. While this assumption did not affect the predicted growth rate (Table 3), we hypothesized the over-estimation of intracellular oxygen consumption could affect metabolic pathway activation and absolute flux values. Additionally, D1 protein repair cost was not included in these previous model simulations. Thus, we simulated photoautotrophic growth in *P. tricornutum* with both the previous 20% assumption and with our experimentally derived Y(II) values and D1 damage rates; comparing the model predictions with experimental O_2_ exchange values derived from membrane inlet mass spectrometry (MIMS).

We defined the model predicted intracellular oxygen consumption as the ratio of the oxygen evolution and the PSII oxygen generation rates. For the LL acclimated condition, both the Y(II) and 20% assumption resulted in almost identical predictions of intracellular O_2_ consumption [24-33% and 24-35% of total PSII O_2_ generation for the Y(II) constrained versus the 20% assumption, respectively; ranges based on Flux Variability Analysis (FVA)]. These values were consistent with previously reported experimentally determined values in LL acclimated *P. tricornutum* of 35±5% (Broddrick *et al*, 2019a). For *P. tricornutum* acclimated to HL, the predicted intracellular oxygen consumption values were dramatically different between the 20% assumption and the Y(II) constrained simulation (63-67% and 40-46% respectively; ranges based on FVA). Using MIMS, we experimentally determined the fraction of PSII O_2_ generation consumed by light-independent mechanisms (maintenance), consumed by light-dependent mechanisms (EET), and evolved (net P_O_) by cells acclimated to high light (600 µmol photons m^−2^ s^−1^). For this HL condition, we determined 42±5% of the PSII generated O_2_ was consumed via intracellular consumption, compared to the Y(II) constrained simulation range of 40-46%. Additionally, the MIMS experiment determined 15±3% of PSII generated O_2_ was consumed via light-independent mechanisms (maintenance), compared to the Y(II) constrained simulation range of 13-14%. Finally, we measured 26±4% of PSII generated O_2_ was consumed via light-dependent mechanisms, compared to the Y(II) constrained simulation range of 26-32%. Overall, while constraining the model with chlorophyll fluorescence data did not affect the growth rate prediction (Table 3), it did result in highly accurate predictions of intracellular oxygen consumption. Table 4 summarizes the Y(II) constrained model predicted photosynthetic parameters.

**Table 4.** Predicted excitation energy flow in *P. tricornutum* acclimated to low and high light. Values are for simulations constrained to account for 100% of absorbed quanta. ΦCO_2_: quantum yield of net carbon fixation, ΦO_2_: quantum yield of net oxygen evolution. Abbreviations: QF: quantum flux, ETR: electron transport rate, PSII: photosystem II, PSI: photosystem I, Y(NO): unregulated excitation energy dissipation, NPQ: non-photochemical quenching.

|  | QF<sup>1</sup> | ETR<sup>2</sup> | Quantum demand<sup>3</sup> | Fraction of QF |  | Charge Separations | ΦCO<sub>2</sub><sup>4</sup> | ΦO<sub>2</sub><sup>4</sup> |
| --- | --- | --- | --- | --- | --- | --- | --- | --- |
|  |  |  |  | PSII | PSI | PSI/PSII |  |  |
| High light | 6.1x10<sup>-10</sup> | 1.4x10<sup>-10</sup> | 16.4 | 0.76 | 0.24 | 0.97 | 0.023 | 0.033 |
| Low light | 1.3x10<sup>-10</sup> | 0.5x10<sup>-10</sup> | 10.5 | 0.60 | 0.40 | 1.05 | 0.044 | 0.066 |

<sup>1</sup> μmol photon cell<sup>-1</sup> s<sup>-1</sup>
<sup>2</sup> μmol electron cell<sup>-1</sup> s<sup>-1</sup>
<sup>3</sup> QF x (0.25 x ETR)<sup>-1</sup>; mol photon x mol<sup>-1</sup> O<sub>2</sub>
<sup>4</sup> μmol x μmol<sup>-1</sup> photon

Next, we investigated how incorporating chlorophyll fluorescence data affected predictions of cross-compartment coupling. Previous modeling in *P. tricornutum* hypothesized excess reductant was shunted from the chloroplast to the mitochondrion (Broddrick *et al*, 2019a), consistent with experimental evidence of energetic coupling of these compartments (Bailleul *et al*, 2015; Murik *et al*, 2019). This previous work suggested photorespiration, branch-chain amino acids, and an ornithine-mediated chloroplast-mitochondrion shunt were the dominant mechanisms for cross-compartment coupling (Broddrick *et al*, 2019a). However, the Y(II) constrained model, validated with the O_2_ values from the MIMS experiment, suggested *P. tricornutum* acclimated to high light has a substantially reduced effective quantum yield at the experimental irradiance [Y(II), Table 2]. Thus, excess excitation energy not performing photochemistry (e.g. NPQ) likely reduces the need for cross-compartment coupling.

We investigated this hypothesis by comparing model predicted intracellular flux using the 20% excitation energy assumption (hereafter 20% assumption) and the combined Y(II) and D1 repair constraints [hereafter Y(II) constrained]. First, we determined the total EET in the system, defined as the excitation energy captured in excess of biomass and maintenance requirements (units: mmol electrons gDW^−1^ h^−1^), as a function of QF for both the 20% assumption and the Y(II) constraints. The results reflected the inaccuracy of the 20% assumption for both LL and HL conditions (Fig. 3A). The biomass normalized EET flux was consistent between LL and HL conditions, and both sets of constraints up to a QF value of approximately 0.3 fmol photons cell^−1^ s^−1^. However, the 20% assumption was a linear constraint applied upstream of both PSII and PSI; thus, the model-predicted EET flux increased linearly with QF. In contrast, EET in the Y(II) constrained model began to plateau (Fig 3), reflecting the decrease in Y(II) and increase in NPQ as QF increased (Fig 2).

**Figure 3.**
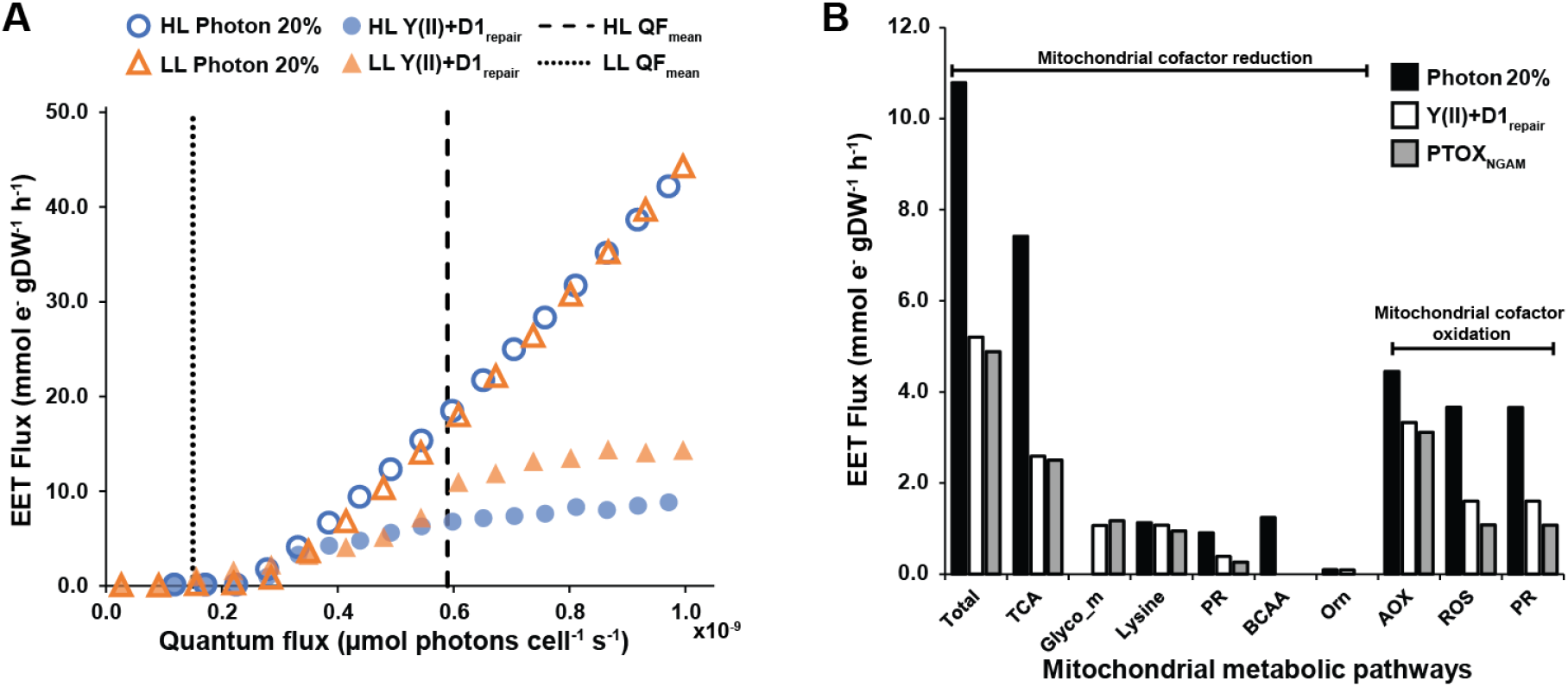
Chlorophyll fluorescence and D1 damage constraints affects model predictions for cross-compartment metabolic coupling. (A) Predicted EET as a function of quantum flux for cells acclimated to LL (triangles) or HL (circles). Open markers: simulations where a fixed 20% of captured photons are lost upstream of the photosystems; filled markers: simulations with Y(II) and D1 repair constraints. Vertical dashed lines represent the mean quantum flux received by the cultures at the experimental irradiance. (B) Total metabolic flux shunted to the mitochondrion via different metabolic pathways for *P. tricornutum* acclimated to high light. Black bars: Simulations where a fixed 20% of captured photons are lost upstream of the photosystems; white bars: simulations with Y(II) and D1 repair constraints; gray bars: simulations with Y(II) and D1 repair constraints and NGAM routed to PTOX. Abbreviations: AOX: alternative oxidase; BCAA: branched-chain amino acid; Glyco_m: mitochondrial glycolysis; Orn: ornithine shunt; PR: photorespiration (reducing: glycine cleavage system; oxidizing: glyoxylate transaminases); PTOX: plastid terminal oxidase; ROS: reactive oxygen species detoxification; TCA: mitochondrial tricarboxylic acid cycle.

Next, we investigated the intracellular EET pathways connecting the plastid and mitochondrion for cells acclimated to HL. The 20% assumption predicted 107% more photosynthetically derived electrons were shuttled to the mitochondria compared to the Y(II) constrained simulation. This difference in cross-compartment energetic coupling resulted in similar cross-compartment shuttles as previously reported (Broddrick *et al*, 2019a); however, the absolute flux through these pathways was altered (Fig. 3B). The Y(II) constrained simulations predicted decreased flux through all pathways, apart from mitochondrial glycolysis, compared to the 20% assumption simulations (Fig 3B). We also explored whether the compartmentalization of light-independent O_2_ consumption [non-growth associated maintenance (NGAM)] affected EET predictions. When we constrained NGAM to the plastid terminal oxidase (PTOX), a thylakoid membrane-localized EET reaction, there was a slight decrease in absolute flux values predicted in mitochondrial EET pathways commensurate with the reduction in photosynthetically derived electrons leaving the plastid ETC. However, the overall trends were consistent with mitochondrial targeted NGAM simulations. An unexplored, potential cross-compartment shuttle suggested by these simulations was the amino acid lysine. However, lysine catabolic flux in the mitochondrion was similar for all conditions (Fig. 3B) suggesting this particular cross-compartment shuttle may not be used as an EET pathway.

Our simulation predictions suggested three routes for plastid-derived reductant consumption: the mitochondrial alternative oxidase (AOX), scavenging of reactive oxygen species (ROS), and conversion of reduced carbon skeletons (glutamate and alanine) to more oxidized forms (alpha ketoglutarate and pyruvate) during glyoxylate transaminase reactions. These last two categories function to detoxify glycolate produced via photorespiration. A unique feature of photorespiration is that it performs both reduction and oxidation of mitochondrial cofactors. Glycine produced by transaminase reactions during the detoxification of glycolate was predicted to be consumed by the glycine cleavage system producing NADH. This reductant then helped fuel the AOX reaction. This linkage between photorespiration, ROS, and AOX is consistent with studies showing AOX to be activated by ROS stress and important in maintaining redox balance in *P. tricornutum* (Murik *et al*, 2019). Our flux predictions come with the same caveat as previous modeling efforts in *P. tricornutum*: experimentally determined intracellular fluxes for this organism have not been adequately determined (e.g., ^13^C metabolic flux analysis) and as such our flux predictions are not yet validated.

### Model-based exploration of bioengineering potential

GEMs account for every known biochemical reaction in the organism and can calculate accurate assessments of resource requirements for biomass components and bioproducts (Dinh *et al*, 2018). We calculated the fraction of linear electron transport (LET) used to biosynthesize each biomass macromolecular fraction (Table 5). The results provided insight into the relative reductant cost of each macromolecular component, which ranged from 177 mmol e^−^ gDW^−1^ of carbohydrate to 411 mmol e^−^ gDW^−1^ of membrane lipids. Using these values and the amount of excess photosynthetically generated reductant in the system, we calculated the theoretical yield of different biomass components if that EET could be rerouted for biomass biosynthesis. The predictions were inversely related to reductant cost with 20.0 mg carbohydrates gDW^−1^ h^−1^ being the highest yield and membrane lipids the lowest at 8.6 mg lipids gDW^−1^ h^−1^ (Table 5).

**Table 5.** Photosynthetically generated electron requirements for different biomass components for cells acclimated to high light. Abbreviations: LET: linear electron flow; EET: alternative electron transport.

| Biomass component | Biomass percent | %LET | Reductant cost<sup>3</sup> | Theoretical yield<sup>4</sup> |
| --- | --- | --- | --- | --- |
| Protein | 70.0 | 43.3 | 304 | 11.6 |
| Structural carb | 6.0 | 2.2 | 177 | 20.0 |
| DNA | 0.3 | 0.2 | 254 | 13.9 |
| Membrane lipids | 3.0 | 2.5 | 411 | 8.6 |
| Pigments | 2.2 | 1.7 | 388 | 9.1 |
| Plastid lipids | 3.7 | 2.6 | 347 | 10.2 |
| RNA | 2.7 | 1.4 | 253 | 14.0 |
| Storage<sup>1</sup> | 10.0 | 5.2 | 258 | 13.7 |
| EET | N/A | 29.1 | N/A | N/A |
| Other<sup>2</sup> | N/A | 11.9 | N/A | N/A |

<sup>1</sup> 2.1 to 1.0 molar ratio of a $\beta$ -1,3-glucan and triacylglycerol (16:1(9Z)/16:1(9Z)/16:0)
<sup>2</sup> Maintenance, vitamins, and cofactors
<sup>3</sup> Millimole photosynthetically generated electrons per gram dry weight of molecule class ( $\text{mmol e}^- \text{gDW}^{-1}$ )
<sup>4</sup> Milligram dry weight component per gram dry weight biomass per hour (mg gDW<sup>-1</sup> h<sup>-1</sup>)

Finally, we explored model-driven engineering strategies to produce high value bioproducts. Bioengineering of photosynthetic microbes has typically targeted fuel or nutraceutical precursors and these targets are normally derived from three different precursors – fatty acids, aromatic amino acids and terpenoids (Brey *et al*, 2020; Kumar *et al*, 2020). We evaluated engineering intracellular pathways to increase flux through plastid fatty acid biosynthesis (hexadecanoate), the shikimate pathway (chorismate), and isoprenoid precursors (isopentenyl pyrophosphate). In our simulation, downregulation of EET provided extra reductant for bioproducts; however, carbon and other elements were also diverted from other biomass components. We simulated diverting up to 50% of cellular biomass to bioproduct synthesis, in increments of 10%, and evaluated changes in the intracellular reaction fluxes that could enable light-drive production of these compounds (Fig 5).

**Fig 5.**
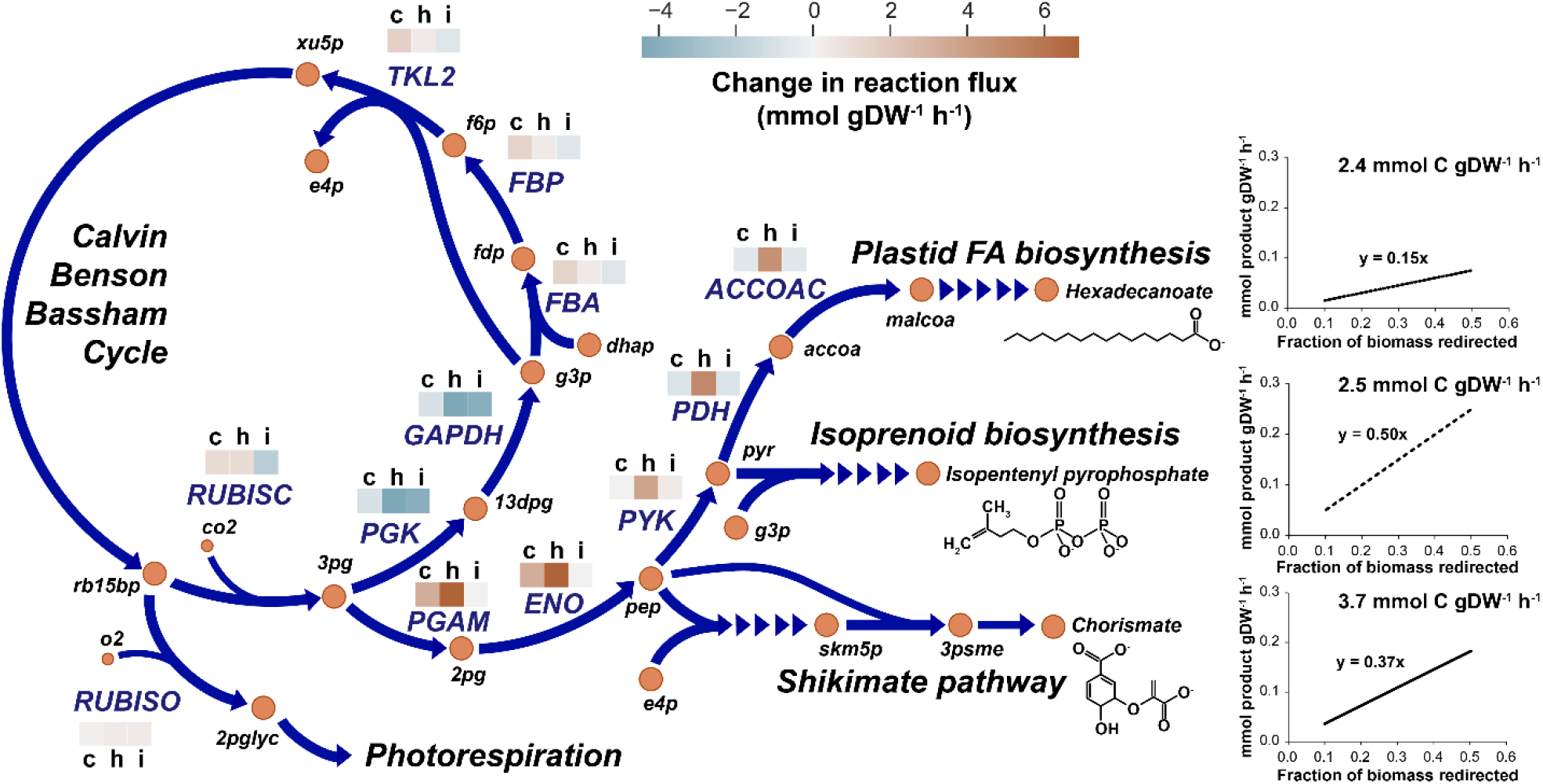
Metabolic engineering potential of *P. tricornutum* acclimated to high light. Changes in metabolic reaction flux towards the bioproducts hexadecanoate, isopentenyl pyrophosphate and chorismate are shown on the flux map. The heatmap above the reactions indicate an increase or decrease in flux towards chorismate (c), hexadecanoate (h), or isopentenyl pyrophosphate (i). Values represent the difference between the baseline simulation fluxes [Y(II) constrained] and bioproduct formation with 30% of biomass rerouted to the desired product. Graphs indicate bioproduct yield as a function of %biomass diverted as well as the carbon-normalized yields. Abbreviations are based on the BiGG Models database (King *et al*, 2016). Abbreviations (Reactions): ACCOAC-acetyl-CoA carboxylase; ENO-enolase; FBA-fructose-1,6-bisphosphate aldolase; FBP-fructose-1,6-bisphosphase; GAPDH-glyceraldehyde-3-phosphate dehydrogenase; PDH-pyruvate dehydrogenase; PGAM-phosphoglycerate mutase; PGK-phosphoglycerate kinase; PYK-pyruvate kinase; RUBISC-ribulose-1,5-bisphophate carboxylase; RUBISO-ribulose-1,5-bisphophate oxygenase; TKL2-transketolase 2. Abbreviations (metabolites): 13dpg-D-glycerate 1,3-diphosphate; 2pg-2-phospho-D-glycerate; 2pglyc-2-phosphoglycolate; 3pg-3-phospho-D-glycerate; 3psme-5-O-(1-carboxyvinyl)-3-phosphoshikimate; accoa-acetyl-CoA; co2-carbon dioxide; dhap-dihydroxyacetone phosphate; e4p-D-erythrose-4-phosphate; f6p-beta-D-fructose 6-bisphosphate; fdp-beta-D-fructose 1,6-bisphosphate; g3p-glyceraldehyde-3-phosphate; malcoa-malonyl-CoA; o2-oxygen; pep-phosphoenolpyruvate; pyr-pyruvate; rb15bp-ribulose-1,5-bisphosphate; skm5p-shikimate-5-phosphate; xu5p-D-xyulose 5-phosphate.

The model predicted a linear increase in product yield as a function of increased biomass diverted to bioproducts. Production rates were 0.15, 0.50, and 0.37 mmol bioproduct gDW^−1^ h^−1^ Fraction_Biomass_^−1^ for hexadecanoate, isopentenyl pyrophosphate, and chorismate, respectively. When normalized to the number of carbons in each of these end products, the yields were 2.4, 2.5, and 3.7 mmol C fixed in bioproduct gDW^−1^ h^−1^ Fraction_Biomass_^−1^ for hexadecanoate, isopentenyl pyrophosphate, and chorismate, respectively. There were no major differences in predicted EET as a result of bioproduct synthesis. We compared the baseline EET [Y(II) constrained simulations above] to the EET of the production strains with 30% of biomass diverted to bioproduct synthesis. The baseline EET was 3.13 mmol e^−^ DW^−1^ h^−1^ compared to 3.08, 3.21, and 3.09 mmol e^−^ gDW^−1^ h^−1^ hexadecanoate, isopentenyl pyrophosphate, and chorismate, respectively. This result suggested the reductant cost of hexadecanoate and chorismate are higher than the mean biomass reductant cost, while the reductant cost of isopentenyl pyrophosphate is lower.

The flux simulations identified metabolic pathways where rerouting of flux is required for bioproduct synthesis. We compared model-predicted metabolic flux routing at an intermediate biomass diversion value of 30%. All three metabolites, hexadecanoate, isopentenyl pyrophosphate, and chorismate, require plastid glycolytic precursors (acetyl-CoA, pyruvate, and phosphoenolpyruvate, respectively). Additionally, chorismate and isopentenyl pyrophosphate both require Calvin-Benson-Bassham Cycle (CBBC) intermediates (D-erythrose-4-phosphate and glyceraldehyde-3-phosphate, respectively) for biosynthesis. These requirements were evident in the reaction flux differences between the reference simulation [Y(II) constrained] and the bioproduct simulations (Fig 5). Hexadecanoate biosynthesis required the largest flux rerouting through lower plastid glycolysis as all the carbon required for its biosynthesis is sourced from acetyl-CoA. For chorismate, six of its ten carbons come from lower glycolysis, requiring increased flux through the reactions phosphoglycerate mutase and enolase. The remaining four carbons are sourced through the CBBC resulting in a slight increase in flux through these reactions, to include a predicted increase in carbon fixation at RUBSICO. While initiation of isopentenyl pyrophosphate biosynthesis utilizes the lower glycolytic metabolite pyruvate, the model predicted the source of this metabolite was recycling of carbon from plastid-mitochondrial metabolic coupling, not redirection of flux away from the CBBC. This result shows how GEMs can result in non-intuitive flux routing towards engineering bioproducts.

## Discussion

In this study, we characterized photoautotrophic metabolism in *P. tricornutum* through integrated chlorophyll fluorescence measurements and genome-scale modeling. Our observations in *P. tricornutum* were consistent with photophysiology under fluctuating and sinusoidal light (Wagner *et al*, 2006) and photoacclimation (Nymark *et al*, 2009). *P. tricornutum* exhibited efficient photoacclimation with the quanta absorbed per pigment remaining consistent between low and high light (Fig. 1B). This efficiency was also observed when looking at the initial slope of the cell-normalized P_O_ versus QF curves (Fig. 1C) and the chlorophyll-normalized P_O_ versus PAR curves (Fig. S1), which were consistent between both low and high light acclimated cultures. This efficiency across a range of photoacclimation conditions likely contributes to the ecological success of diatoms in dynamic light environments (Behrenfeld *et al*, 2021).

Our interpretation of photophysiology was heavily influenced by analyzing the P_o_ and PAM data with quantum flux (QF) as the independent variable. The 1-qL versus QF curves (Fig. 1D), the shape of the chlorophyll fluorescence parameters versus QF curves (Fig. 1 E, F), and the D1 content as a fraction of total protein (Table S1) were consistent between low and high light. Contrasting PAM vs. QF (Fig. 1E, F) with PAM vs. PAR (Fig. S5A, B) illustrates how interpreting photophysiology from a QF perspective affects conclusions about photophysiology. Using QF as the independent variable not only normalized the comparisons across light regimes (fluorescent bulb for culturing, red LED for PAM experiments) and experimental apparatus (e.g., Roux flasks for culturing versus a round cuvette for PAM measurements), this approach also accounted for inherent changes in photophysiology (e.g., pigment content, photochemical efficiency).

We observed very little dissipation of excitation energy via NPQ at the experimental QF values (Fig. 1E, F). For the HL conditions, this lack of NPQ was coupled with an effective quantum yield of PSII [Y(II)] value of 0.32 (Table 2) suggesting the presence of alternative dissipative mechanisms [Y(NO)]. NPQ has been shown to be an important excitation energy dissipation mechanism in dynamic light conditions (Olaizola *et al*, 1994; Lavaud *et al*, 2002; Wagner *et al*, 2006); however, our results suggest these other dissipative mechanisms are sufficient to prevent photoinhibition under stable light environments. Overall, these data suggest *P. tricornutum* employs a photoacclimation strategy that emphasizes rapid utilization and dissipation of light energy. This strategy results in conditions where the overall photosynthetic apparatus is under-utilized, as in our low light acclimated cultures. However, our rapid-light-curve experiments suggest this allows *P. tricornutum* to immediately respond to an increase in available photon flux without the need to biosynthesize additional macromolecules, evident from consistency in total photosystem II content per cell and the redox state of the plastoquinone pool as a function of quantum flux (Fig. S2A, B).

Translating the QF to a photon uptake constraint and P_O_ into an oxygen evolution constraint in the genome-scale model resulted in accurate predictions of photoautotrophic growth (Fig. 6). Growth rate in the HL condition was underestimated by 12%; however, this was likely due to the beginning of carbon limitation in the sample during short-term measurements of photosynthetic capacity (Fig. S3A). We chose not to spike in exogenous bicarbonate for our O_2_-evolution measurements as we were interested in measuring photosynthetic parameters relevant to our culturing conditions. However, our simulations underestimated the growth rate at high light, suggesting the carbon environment needs be considered when performing rapid light curves (indeed, standard protocol is to include several mM NaHCO_3_^−^ in these assays to avoid carbon limitation). Our efforts to establish the hierarchy of constraints suggested photon uptake and chlorophyll fluorescence constraints alone accurately predict growth rate at low acclimation irradiances (Table 3). This opens the possibility for non-invasive monitoring of culture health and productivity for biotechnology applications. Bypassing a portion of a production culture through a passive sampling window with integrated chlorophyll fluorescence and spectral absorption analysis, coupled with data on surface irradiance, would recreate the chlorophyll fluorescence and photon uptake constraints. As bioproduction conditions are usually high density, which would likely result in low cell-specific quantum flux, integrating non-invasive sampling with our modeling construct could result in accurate measurements of photoautotrophic metabolism in these settings.

Constraining biomass accumulation with QF and P_O_ automatically predicts relevant photosynthetic parameters in a manner similar to previous investigations (Wagner *et al*, 2006; Jakob *et al*, 2007). However, GEMs also predict the optimal distribution of excitation energy between PSI and PSII, an advantage compared to previous work where it was assumed 50% of the absorbed quanta were directed to each photosystem. Our simulations predicted a two-fold increase excitation energy was utilized by PSII under high light conditions compared to low light conditions, resulting in approximately 76% of absorbed quanta directed to PSII. However, there was a similar number of charge separation events at both photosystems (Table 4). This is consistent with the observation that there is minimal cyclic electron flow (CEF) around PSI in *P. tricornutum* (Bailleul *et al*, 2015) and energy dissipation of light energy at PSII Y(NO), which would result in roughly equivalent charge separations at both photosystems. Our model derived ETR differs from methodologies that assume equal excitation energy routed to both photosystems and a lower predicted quantum demand at experimental QF values (Table 4). For an organism like *P. tricornutum* that does not employ extensive CEF, the advantage of this approach is diminished. However, for microalgae that dynamically reroute excitation energy between the photosystems and adjust their biomass macromolecular composition as part of their photoacclimation strategy [e.g. *Chlamydomonas reinhardtii* (Davis *et al*, 2013; Lucker & Kramer, 2013)], implicitly calculating excitation routing between photosystems using our modeling framework will result in better approximations of ETR.

Incorporation of chlorophyll fluorescence measurements as a GEM constraint increased the accuracy of model predictions. Accounting for photon loss upstream of the photosystems resulted in a more accurate prediction of intracellular oxygen production and reductant-mediated oxygen consumption. These new constraints affected predictions related to excess reductant in the system (EET) and cross-compartment metabolic coupling (Fig. 3A, B). The plateau in EET flux as a function of QF (Fig. 3A) was correlated to NPQ activation (Fig. 1F), suggesting saturation of EET pathways triggers this photoprotective mechanism. Our results suggest the following model of photophysiology in *P. tricornutum* in a stable light environment: up to a QF of approximately 0.3 fmol photons cell^−1^ s^−1^, Y(NO) dissipates excess excitation energy until the steady-state reduction of the plastoquinone pool is approximately 50%. At this point, EET is activated to facilitate re-oxidation of the photosynthetic electron transport chain. At a QF of approximately 0.6 fmol photons cell^−1^ s^−1^, the EET pathways are saturated, the steady-state reduction of the plastoquinone pool reaches 70%, and NPQ is activated to assist in dissipating captured photons. Therefore, the model accurately recreates the onset of NPQ that occurs when light absorption outpaces the ability to utilize this energy within metabolism. Interestingly, this model is consistent for cells acclimated to both low and high light, and *Phaeodactylum* is known to maintain high capacity for NPQ in both light conditions (Taddei *et al*, 2018).

Currently, experimentally derived photoautotrophic metabolic flux values for *P. tricornutum* do not exist, thus our flux predictions are hypotheses that still require validation. Still, the approach outlined in this study is generally applicable to all phototrophic genome-scale model simulations and previous efforts using experimentally-derived electron transport efficiencies, as opposed to PAM, showed good agreement with ^13^C metabolic flux analysis (Broddrick *et al*, 2019b).

The interest in integrating light-driven metabolism into bioengineering and synthetic biology necessitates an iterative framework for bioprocess development. The design-build-test-learn (DBTL) paradigm is one such approach that has been leveraged to rapidly increase bioproduct titers (Petzold *et al*, 2015; Carbonell *et al*, 2018). Computational tools are integral to these workflows. Relevant to the *design* step of the process, GEMs have been extensively used to rationally engineer metabolism to generate a wide variety of phenotypes (Czajka *et al*, 2021; Bang *et al*, 2020; Li *et al*, 2019). We explored implementing the modeling framework towards light-driven bioproduct formation, as adoption of these approaches is underrepresented in phototrophic systems. Initial assessments quantified the reductant cost of cellular macromolecules, which provided insight into the theoretical yield of different compound classes (Table 5), with protein biosynthesis being the predominant sink of photosynthetic energy. EET was found to consume 29% of LET in high light. These EET pathways serve to oxidize the photosynthetic electron transport chain and resupply low energy cofactors to autotrophic metabolism. However, they are generally viewed as wasteful as this energy could be utilized to increase biomass yields (Peers, 2014). We utilized our model to estimate the metabolic consequences of redirecting metabolism towards important chemical precursors. The overexpression of plastid fatty acids (hexadecanoate), the shikimate pathway (chorismate), and isoprenoid precursors (isopentenyl pyrophosphate) were only partially fueled by reductant that would normally be dissipated by EET, showing there is still considerable potential associated with the engineering of primary photosynthetic metabolism to increase the overall yields of bioproducts.

Furthermore, our modelling suggests that increasing the relative flux of reduced carbon to metabolic precursors of industrial interest may require significant engineering of central carbon metabolism. For instance, increasing the production of our three selected metabolites increased the flux of 3-phospho-D-glycerate (3pg) through the plastid glycolytic pathway (Figure 5). Additionally, the increased flux of carbon to isoprenoid biosynthesis or through the shikimate pathway puts an increased demand for biosynthetic intermediates that are sourced from the CBBC (glyceralde-3-phosphate and D-erythrose-4-phosphate, respectively). Photosynthetic microorganisms, which include diatoms, naturally redirect carbon and energy to storage molecules under conditions of nutrient deprivation, such as nitrogen limitation (Alipanah *et al*, 2015). This phenotype forms the basis for much of the interest in biofuel applications of these phototrophs (Levering *et al*, 2015). Thus, a reasonable strategy and potential future direction is the diel separation of carbon fixation and bioproduct formation. It may be that rerouting these metabolites from sugar polymer degradation via the pentose phosphate pathway or mitochondrial β-oxidation of lipids (Jallet *et al*, 2020) may alleviate some of the pressure on the CBBC.

The next step is to build and test these strains to initiate the first iteration of the DBTL cycle. During the *test* phase, the model constraints provide a roadmap for relevant process parameters and a framework to evaluate process performance, to include assessments on the efficiency of EET usage. It is important to note our design simulations do not include possible changes in photophysiology due to strain engineering (e.g., an increase in P_O_). However, physiological outputs from the testing of the strain designs proposed can be re-integrated into the modeling framework described here to include changes in experimentally derived constraints. This contributes to the *learn* step of the DBTL cycle, closing the loop on the first iteration and enabling an updated design strategy for the next iteration.

Taken together, our results show integrating relevant measurements of photosynthetic physiology with genome-scale models results in quantitative predictions of condition-specific phenotypes. This paves the way for iterative design and real-time process control of photobioproduction platforms.

## Methods

### Culture conditions

*Phaeodactylum tricornutum* CCAP 1055/1 was grown axenically in silicon-free Instant Ocean artificial seawater (salinity 35‰). Nutrients were added according to the stoichiometry described in (Guillard, 1975) but at 2.3 fold higher concentrations to avoid nutrient limitation. Cells were cultured at 18°C in 400 mL medium in 1 L Roux flasks. Flasks were bubbled with air (1 L air L^−1^ culture minute^−1^) under continuous illumination in a temperature-controlled incubator. Cultures were light acclimated (low light (n=3) at 60 µmol photons m^−2^ s^−1^, high light (n=4) at 600 µmol photons m^−2^ s^−1^) for 72 hours, diluted and grown until mid-exponential phase before being harvested.

### Cell physiology measurements

Cell densities were determined using a BD Accuri C6 flow cytometer as described in (Jallet *et al*, 2016a). Growth rates were determined based on the change in cell counts from inoculation to harvest. Cell dry weight was determined by taking 50 mL of culture (n=3) and filtering onto a GF/C glass microfiber filter (diameter: 47mm). Filters containing cellular biomass and media controls (n=3) were dried at 95°C overnight. Cellular dry weight was determined by subtracting the post-drying mass from the pre-drying mass, after normalizing to the media control.

### Determination of cell dimensions

One µL of Lugol’s solution was added to 1 mL of culture and the sample was stored at 4°C until analysis. For imaging, thin pads of 1% (wt/vol) agarose were prepared using Mini-PROTEAN R Tetra Cell Casting Module. From this gel, 1-2 cm square pads were cut and placed onto a microscope slide and 2-5 µl cell culture liquid was added to the pad and let dry. Then a microscope slide cover was gently placed onto of the agarose pad and cells were imaged using a DeltaVision inverted epi-fluorescence microscope (Applied Precision, Issaquah, WA). Images were captured using a CoolSnap HD charge-coupled device (CCD) camera (Photometrics, Tucson, AZ). Cell length and width were determined using the straight-line tool in ImageJ (Schindelin *et al*, 2015) and used to determine cell volume [high light (n=94) and low light (n=46) acclimated cells]. *P. tricornutum* was modeled as a core ellipse with two cones extending away from the core ellipse. The core ellipse was calculated according to the following equations:

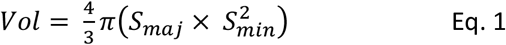

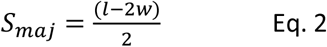

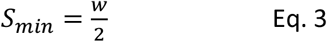

where l is the cell length and w is the cell width.

The cones were calculated according to the following equation:

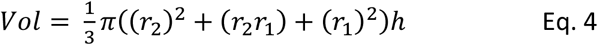

where r_2_ is equal to w/4, r_1_ is equal to 1 µm and h the cell width. Mean and standard deviation were determined using the Python Numpy package (Harris *et al*, 2020).

### Pigment extraction

Cells (4 mL of liquid culture) were collected by centrifugation at 10,000 x g at 5°C for 15 minutes. The supernatant was discarded and the cell pellet was frozen at −80°C until processing. Chlorophyll was extracted with 50 µL DMSO and 1950 µL of methanol, incubated in the dark for 30 minutes, and centrifuged at 10,000 x g at 5°C for 15 minutes. The pigment containing supernatant was transferred to a 1 cm path length cuvette. Absorbance spectra were collected using a Cary 60 UV-Vis Agilent spectrophotometer in scan mode (350-800 nm, scan interval of 1 nm). Chlorophyll *a* and *c* concentrations were determined using the equations for methanol (Ritchie, 2008).

### Cellular absorption coefficients

Cellular absorption coefficients were determined based on published protocols (Moore *et al*, 1995). Approximately 5×10^7^ cells were collected by filtration onto a GF/A glass microfiber filter (21 mm diameter). The filter was cut to fit in a 1 cm path length cuvette and placed against the inside of the cuvette. Absorbance spectra were collected used a Cary 60 UV-Vis Agilent spectrophotometer in scan mode (350-800 nm, scan interval of 1 nm). A total of 4 technical replicates were collected per sample and averaged. To decrease the noise from filter scattering, the absorbance spectra were smoothed using a Savitzky-Golay filter (width= 17 nm, polynomial = 2nd order) as implemented in SciPy (Virtanen *et al*, 2020). The resulting spectra were blank subtracted against the appropriate media and normalized to an OD_750_ value of 0. The wavelength specific absorption coefficient was determined, along with correcting for filter amplification according to the following equation:

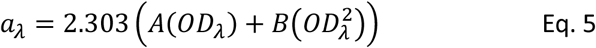

where OD_λ_ is the absorbance at a given wavelength and A and B are species-specific coefficients for the β-factor correction (0.388 and 0.616) (Finkel & Irwin, 2001). The cell normalized absorption coefficient (a*_cell_, units: cm^2^ cell^−1^) and the pigment normalized coefficient (a*_pigm_, units: cm^2^ µg^−1^ pigments) were determined by dividing a_λ_ by either the total number of cells deposited on the filter or the total pigment mass, respectively, and then multiplying the resulting value by the filter area onto which the cells were deposited (2.1 cm^2^ for the 21 mm diameter GF/A filter).

### Simultaneous oxygen evolution and chlorophyll fluorescence parameters

Rapid light curves (RLCs) were performed as outlined previously (Jallet *et al*, 2016a; Broddrick *et al*, 2019b). A Walz Dual PAM 100 fluorometer in a temperature controlled custom cuvette holder and a FireSting Optical Oxygen Meter were used for the simultaneous measurement of chlorophyll fluorescence and oxygen evolution. Approximately 30 mL of culture was removed, and cells were pelleted by centrifugation (3000 x g, 10 minutes at the experimental temperature). Cell pellets were resuspended in fresh media to the target cell density (HL: 2×10^7^ cells mL^−1^, LL: 1×10^7^ cells mL^−1^) and kept in the dark for 10 minutes prior to analysis. For select experiments, the cells were reconstituted in fresh media supplemented with 5 mM sodium bicarbonate. Dark respiration rates were collected for approximately 10 minutes prior to running RLCs. A red actinic light (635 nm) was used to provide a saturating pulse (600 ms, 10,000 µmol photons m^−2^ s^−1^) for fluorescence measurements. Cells were step illuminated (HL: 60 seconds, LL: 90 seconds) at the following increasing intensities (µmol photons m^−2^ s^−1^):

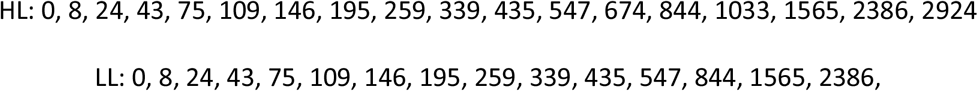

followed by the red actinic saturating pulse.

The chlorophyll fluorescence parameters F_v_/F_m_, Y(II), 1-qL and NPQ were determined as described (Schreiber *et al*, 1995; Kramer *et al*, 2004). Shading in the round cuvette was accounted for by calculating the attenuation across the cuvette path length as described previously (Broddrick *et al*, 2019b):

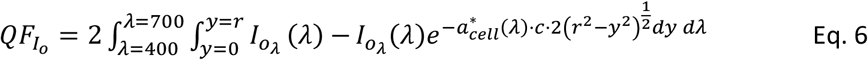

where QF_Io_ is the quantum flux in µmol photons m^−2^ s^−1^ at a given PAR value (I_o_), λ is the wavelength, Io_λ_ (λ) is the fraction of the PAR at a given wavelength λ, r is the radius of the cuvette (0.56 cm), a*_cell_ is the wavelength-specific absorption coefficient in cm^2^ cell^−1^, and c is the cell density in cells cm^−3^. QF was converted to µmol photons cell^−1^ s^−1^ by multiplying QF_Io_ by the rectangular surface area of the cuvette (width = 0.56 cm, height = 1.15 cm), converted to m^2^ and divided by the total number of cells in the cuvette. This QF value was used as the independent variable in plots of oxygen-based photosynthesis (P_O_) versus QF.

Time-course measurements of oxygen evolution were exported from the FireSting O2 Logger software as a .txt file. The data was aligned to the PAM data irradiance values and the first 10 seconds of data after each increase in irradiance was discarded as the O_2_ evolution rate stabilized at the new irradiance value. The oxygen evolution rate was determined by taking the slope of the O_2_ versus time plot for the remaining time interval (50 seconds for HL, 80 seconds for LL) using the Python SciPy package (scipy.stats.linregress) (Virtanen *et al*, 2020). The resulting oxygen evolution rates were then normalized to cell counts. Dark period respiration rates were determined in a similar manner by taking the slope of the oxygen consumption versus time curve for the last 2 minutes of the 10-minute dark acclimation period.

### Membrane inlet mass spectrometry (MIMS)

Membrane inlet mass spectrometry was measured similarly to (Broddrick *et al*, 2019a) and (Ware *et al*, 2020). Cells were collected for chlorophyll quantification according to (Jallet *et al*, 2016a), with spectrophotometric quantification performed in 100% methanol according to the equations provided in (Ritchie, 2008). Samples corresponding to 4 µg chlorophyll *a* mL^−1^ were collected and resuspended in fresh f/2 media. All steps were performed at 18°C. 2 mL of culture was loaded into a custom made cylindrical Accura ClearVue SLA cuvette. Oxygen evolution and consumption was measured using a quad mass spectrometer (Pfeiffer PrismaPlus QMG220, Quadera v4.62). Dissolved gas was pulled through a silicon membrane, connected to the mass spectrometer via a stainless-steel tube with a vapor trap (ethanol and dry ice). Cells were bubbled with N_2_ to deplete oxygen (^16^O_2_, m/z = 32) from the suspension to approximately 50% of atmospheric oxygen concentrations. Cells were kept under constant stirring via a magnetic stir bar. The suspensions were injected with ^18^O_2_ (Cat #490474, Aldrich) and mixed for 10-15 minutes until equilibration was achieved. ^18^O_2_ was then purged from the sample using a stopper leaving a 1.4 mL final culture volume. Gas consumption was measured in the dark for 5 minutes to calculate the respiration rate. Cells were then illuminated with a blue measuring light (Walz, Dual-PAM-100) to achieve a 0.2V fluorescence signal for 15 minutes to relax photoprotective processes. A white LED programmed to achieve 600 or 60 µmol photons m^−2^ s^−1^ (measured with Walz, ULM-500, US-SQS/L attachment) was used to illuminate cultures to their corresponding in situ light intensity for 7 minutes. The slopes of oxygen consumption (^16^O_2_, m/z = 32, and ^18^O_2_, m/z = 36) and evolution (^16^O_2_, m/z = 32) were calculated according to (Beckmann *et al*, 2009) using the mass charge (m/z) change on a per second basis, calculated over the last 30s of illumination. Argon (m/z = 40) was used to normalize oxygen concentrations, minimizing the effects of pressure change and abiotic gas consumption (Bailleul *et al*, 2015).

### Genome-scale metabolic modeling

#### Model and constraints

We used the *P. tricornutum* genome-scale model (GEM) iLB1034 (Broddrick *et al*, 2019a). Simulations were performed in a similar manner to (Broddrick *et al*, 2019b). The biomass objective function was updated to account for differences in pigments between the low and high light conditions (Table 1). Photoautotrophic growth was simulated for a 12-hour (high light) or 24-hour (low light) growth period broken into 20 minute pseudo-steady-state segments. Light was modeled coming from the side of the flask. The Roux flasks had approximately 375 mL of culture at the time of the experiments resulting in a light-facing surface area of 80 cm^2^ and a path length of 4.7 cm. At the beginning of each simulation, the appropriate constraints were updated. First, the total biomass in the culture was divided by the cell dry weight to determine the total cells in the culture. Next, the photon uptake rate was determined by calculating the total light absorption along the 4.7 cm path length. We used the spectral distribution of photon flux for the given light source at the experimental irradiance (I_o_(λ)), the cell specific spectral absorption coefficient (a*_λ_), and the cell density (cells mL^−1^), to determine the photon uptake flux (I_a_) in units of µmol photons (time interval)^−1^ using the following equation:

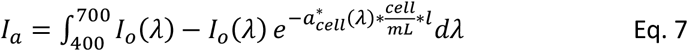

The sum of absorbed light at each wavelength between 400 and 700 nm was used to set the reaction bounds of the photon exchange reactions in the GEM (reaction ID: EX_photon_e).

The P_O_ vs. QF curves were fit to a Platt [128] equation for photosynthesis prediction (P), using quantum flux as the independent variable.

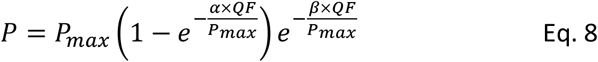

P_max_ is the maximum photosynthetic rate, and α and β are parameters that describe the initial slope of the curve, and the photoinhibition (if present), respectively. The respiration rate was added to all values prior to generating the fit since the Platt curve is forced through the origin. The respiration rate was then subtracted from the fit to return the curve to the gross oxygen evolution rate. These curves were used to determine the oxygen evolution rate based on the total absorbed quantum flux and used to set the bounds of the oxygen exchange reaction in the GEM (reaction ID: EX_o2_e).

#### Integrating chlorophyll fluorescence measurements into the genome-scale model

Chlorophyll fluorescence parameters were incorporated into the GEM based on the experimental values for Y(II), Y(NPQ) and Y(NO). The Y(II) vs. QF data was fit to an exponential decay function 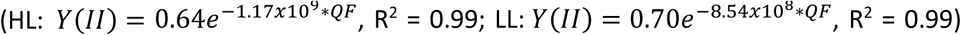. Y(NPQ) vs. QF was fit to a Hill function:

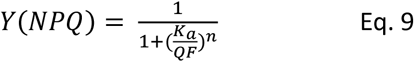

Where HL: n=5.5, K_a_ = 1.37 × 10^−9^ and LL: n=3, K_a_=2.30 × 10^−9^. For each simulation, the calculated QF was used to determine the Y(II) and Y(NPQ) values. Y(NO) was defined as:

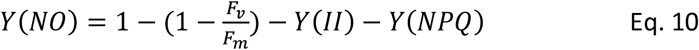

The fraction of absorbed photons available to perform photochemistry was constrained using the following model reaction:

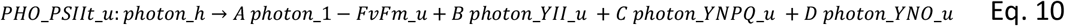

Where PHO_PSIIt_u is the model reaction name, photon_h is the pool of excitation energy available to both PSII and PSI to perform photochemistry (equivalent to quantum flux, QF), A is the fraction of excitation energy lost upstream of the photosystems, B is the Y(II) value at the experimental QF, C is the Y(NPQ) at the experimental QF, and D is Y(NO) at the experimental QF. Only photon_YII_u was included in the model PSII reaction; thus, limiting the amount of excitation energy to perform photochemistry to the Y(II) fraction.

#### Non-growth associated maintenance (NGAM)

NGAM was calculated from the experimental dark respiration rate. This value was set as the lower bound for a fictional quinone oxidase (reaction ID: NGAM), which forces a minimal amount of reductant mediated oxygen consumption consistent with the observed dark respiration rate.

#### Determination and integration of damage to the D1 subunit of PSII

Cells were cultured as outlined above. Exponential phase cultures were split into control and lincomycin treatments in biological triplicates. Lincomycin infiltration was achieved by inoculating cultures with 500 µg mL^−1^ lincomycin for 10 minutes in the dark (Key *et al*, 2010). Samples were collected for cell counts, absorption spectra, F_v_/F_m_ and Western blots at 0, 15, 30, 60 and 90 minutes. 2 mL of cells were collected in Eppendorf tubes and transferred to 10 μmol photons m^*−*2^ s^*−*1^ with a red-light filter for 20 minutes to relax NPQ. Cells were collected on a glass fiber prefilter (Merck Millipore Ltd). A measuring light intensity was applied to elicit a 0.15-0.2 V instrument response at the experimental irradiance. It was provided for 30 sec for accurate determination of F_0_. A saturating pulse (600 ms, 10,000 μmol photons m^*−*2^ s^*−*1^, 635 nm) was applied to determine F_m_, and calculate the maximum photochemical quantum yield of PSII (F_v_/F_m_). The D1 concentration as a fraction of total protein was determined by SDS-PAGE and Western blots as detailed in (Jallet *et al*, 2016a) with some minor modifications. For Western blot analysis, 0.8 µg of total sample protein in 25 µl total volume was loaded into Novex WedgeWell 10-20% Tris-Glycine Gels. Alongside samples, PsbA D1 protein quantitation standards (Agrisera, #AS01 016S) were run. Standards were loaded at the same volume as samples, being diluted in the same sample buffer (Jallet *et al*, 2016a). D1 protein standards were loaded at three concentrations, ensuring samples fell within linear range. Standards and samples D1 protein content was quantified by densitometry (ImageJ, v1.53 https://imagej.nih.gov/ij/download.html). Values from the quantification of sample D1 protein concentration were divided by total protein loaded to calculate D1 protein as a percentage of total protein.

The D1 decay rate in lincomycin treated cells was determined using equation 5 from (Campbell & Tyystjärvi, 2012).

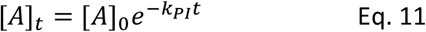

The first-order decay constant, k_PI_ was calculated from the from the change in the maximum quantum yield of PSII (F_v_/F_m_) between cells with and without lincomycin treatment. [A]_0_ was determined from the D1 content as a fraction of total protein calculated from the Western blot analysis. The steady-state D1 damage rate in percent total protein h^−1^ is equal to the first derivative of the above equation solved at a given steady-state fraction of D1 subunits undergoing repair:

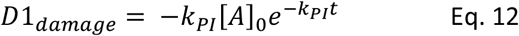

We approximated the steady-state fraction undergoing repair using the difference between F_v_/F_m_ and Y(II) values in the RLCs with a 10 s blue light treatment, at the experimental QF (Fig. S4), under the hypothesis the residual decrease in quantum efficiency was due to inactive PSII complexes undergoing repair. The final D1 damage constraint in mmol D1 gDW^−1^ h^−1^ was calculated with the following equation:

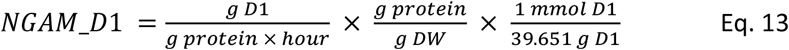

Where the first term is the steady-state damage rate, the second term is the protein fraction of the total biomass (0.70 for high light, 0.64 for low light) and the third term is the formula mass of the *P. tricornutum* D1 subunit (39,651 Da). The model constraint was set from the mean steady-state damage rate and the upper and lower bounds were calculated by combining the variances of ±1 percent standard deviation of the Western blot analysis, ±1 percent standard deviation of the k_PI_ value from the lincomycin treatment data, and a 5% error on the biomass protein fraction for a total of 20% standard deviation for the high light value and a 29% standard deviation for the low light value.

#### Genome-scale model simulations

The simulation was performed by maximizing the biomass objective function (BOF) reaction using the parsimonious FBA function (Lewis *et al*, 2010) as implemented in COBRApy (Ebrahim *et al*, 2013). The flux through this reaction is equal to the biomass accumulation in milligrams over the 20-minute time interval. This biomass was added to a running total of the total culture biomass and used to parameterize the next 20-minute simulation interval. The simulations included the mean and ±1 standard deviation of the a*_cell_ values, cell dry weight to determine the cell count at each time interval, and the oxygen evolution rate versus QF. This resulted in a set of 27 parameters for which growth rate was determined using the following equation:

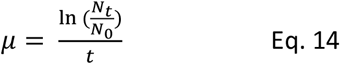

The mean, maximum and minimum growth rates are reported. For the bicarbonate spiked simulation, the mean values were used for all parameters except the oxygen uptake rate which was parameterized using a Platt fit through the bicarbonate spiked P_O_ vs. QF curve. Reaction fluxes were exported into a csv file with Python and visualized in Escher (King *et al*, 2015).

For simulations with varying constraints, the model was parameterized with a combination of the following constraints (unconstrained refers to an arbitrarily high value that doesn’t limit growth): photons (*hv*): the model photon exchange reaction (EX_photon_e) was set to the calculated QF value; oxygen evolution (P_O_): the model reaction EX_o2_e lower bound was set to the experimental oxygen evolution rate at the calculated QF; D1: the D1 damage rate was included; Y(II): the pseudo-reaction PHO_PSIIt_u that accounts for the 1-F_v_/F_m_, Y(II), Y(NPQ) and Y(NO) fractions was included. To calculate excess energy in the system a demand reaction was added to the model upstream of the photosystems that allowed any excess QF to leave the model (reaction ID: DM_photon_c). The flux through this reaction is equal to the excitation energy in excess of the requirements to generate biomass and satisfy NGAM (see above). For all other simulations, the bounds of this reaction were set to 0. For analyses that required a range of feasible fluxes through a model reaction, flux variability analysis (FVA) was used as implemented in COBRApy (Ebrahim *et al*, 2013) with the ‘loopless’ option set to ‘True’.

#### Bioengineering applications simulations

Linear electron transport (LET) was defined as the flux through photosystem I (PSI) minus the flux through cyclic electron flow (CEF). To determine the %LET allocated to each biomass component, we first determined the EET in the system by opening the DM_photon_c reaction, set the lower bound of biomass objective function (BOF) to 99.9% of maximum and then maximized the DM_photon_c flux. This set the baseline LET required for generating biomass. We then iterated over each biomass component, removing the component from the BOF and then adjusting the BOF reaction bounds to (1-biomass component fraction) * maximum. Flux through the DM_photon_c reaction was once again maximized and the difference between this flux value and that for the full BOF was considered the photons necessary to produce the removed biomass component.

Bioproduction of representative compounds were performed by selecting a model metabolite for biosynthesis (plastid fatty acids: hexadecanoate, model id: hdca_h; the shikimate pathway: chorismate, model id: chor_h; and isoprenoid precursors: isopentenyl pyrophosphate, model id: ipdp_h). The default model was solved for maximum biomass (in mg cell dry weight), and subsequent simulations fixed the biomass production at 10% intervals from 50% to 90% of this maximum. A demand reaction was added to the model allowing the representative pathway metabolite (hdca_h, chor_h, or ipdp_h) to leave the system. The model objective was set to maximize this demand reaction. EET was determined as outlined above for all combinations of biomass reallocation and metabolite production. All calculations and simulations were performed using in-house scripts developed in IPython (Fernando *et al*, 2007). For pathway engineering analysis, the reaction fluxes for the 70% biomass results (30% of cellular biomass re-routed to bioproduct formation) were exported into a csv file with Python and visualized in Escher (King *et al*, 2015). All simulation code, models, flux simulations and metabolic maps are available in the Supplemental Material.

## Data Availability

All code used to analyze and generate the results and figures for this study, along with input data, can be found at https://github.com/JaredTBrod/PAM_GEMs.

## Acknowledgements

The authors would like to thank Dr. David G. Welkie and Prof. Susan S. Golden at UC San Diego for their assistance in imaging *P. tricornutum* for the cell size measurements, as well as Marc Abrams for his critical review of the manuscript.

## Supplemental Figures and Tables

**Table S1:** Photosystem II D1 subunit damage rates and resulting maintenance cost for *P. tricornutum* acclimated to high and low light.

|  | D1 damage constant<br>(min<sup>-1</sup>) | D1 protein<br>(% total protein) | D1 damage constraint*<br>(mmol D1 gDW<sup>-1</sup> h<sup>-1</sup>) | Maintenance cost<br>(mmol NTP gDW<sup>-1</sup> h<sup>-1</sup>) |
| --- | --- | --- | --- | --- |
| Low Light | 6±1 x10<sup>-4</sup> | 1.1±0.2 | 7±2 x10<sup>-6</sup> | 8±2 x10<sup>-3</sup> |
| High Light | 2.33±0.17 x10<sup>-2</sup> | 1.1±0.1 | 2.53±0.51 x10<sup>-4</sup> | 0.27±0.06 |

\* Protein fraction in the model biomass function is 0.70 for HL and 0.64 for LL.

**Figure S1.**
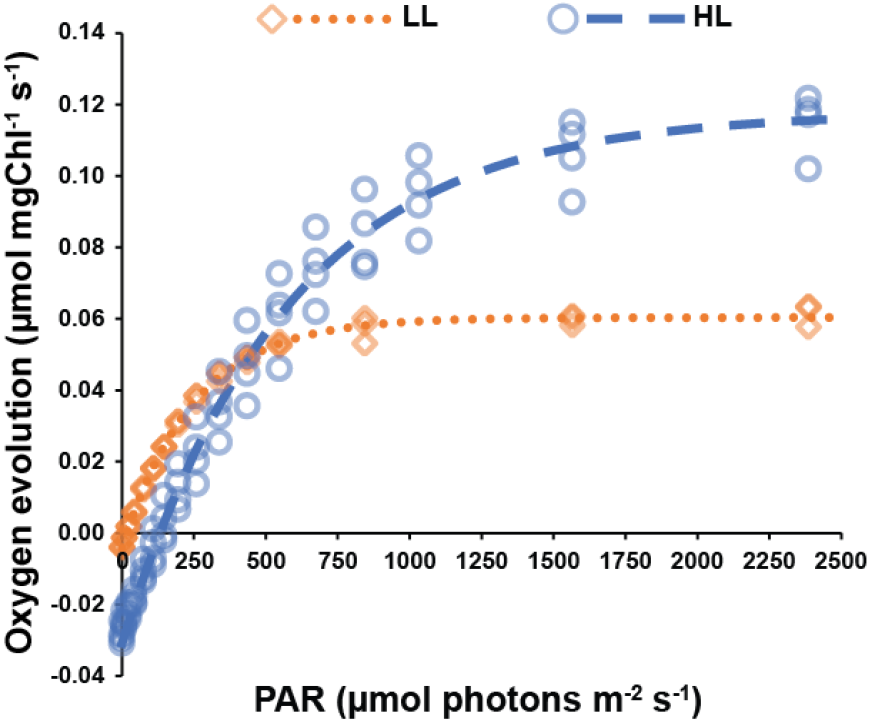
Chlorophyll-normalized photosynthetic rate versus quantum flux. Chlorophyll normalized P_O_ versus PAR curve. Abbreviations and definitions: LL: low light, HL: high light, PAR: photosynthetically available radiation. Data based on n=3 biological replicates for LL and n=4 biological replicates for HL.

**Figure S2.**
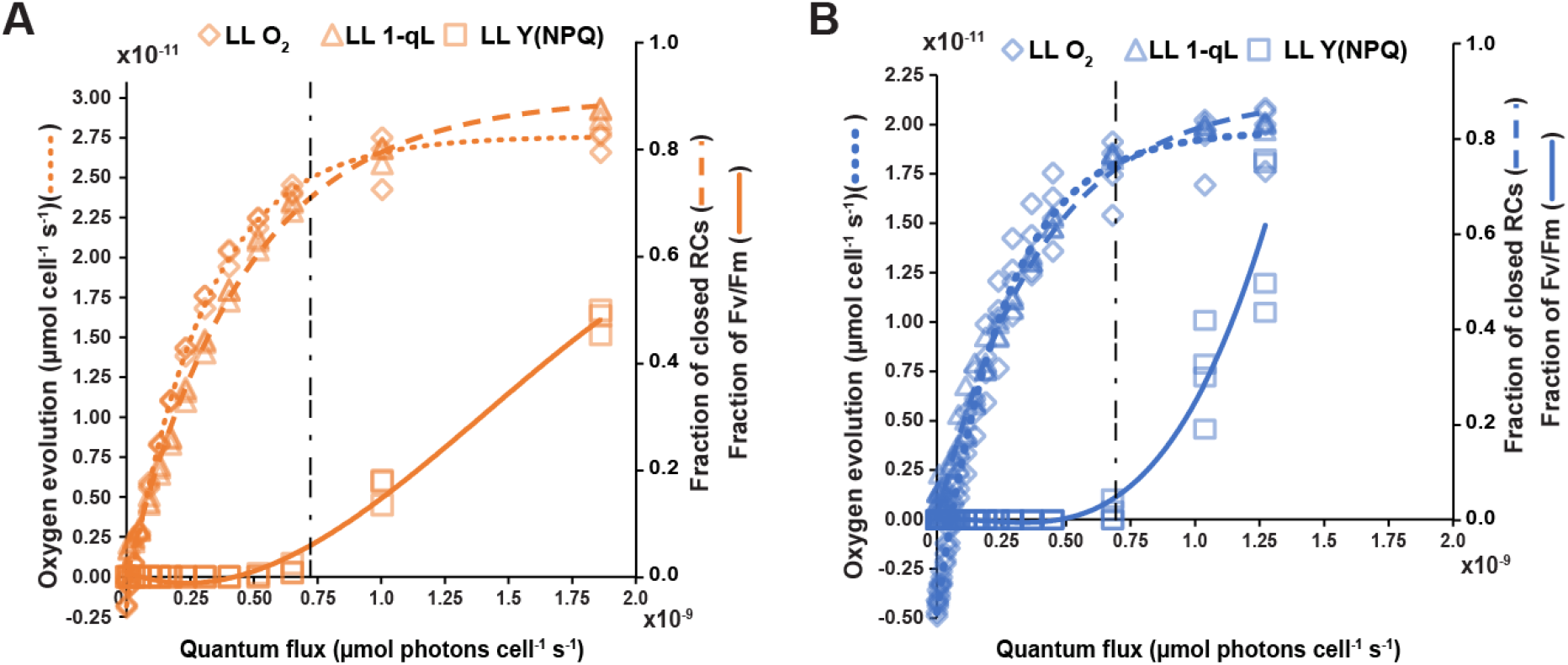
Correlation between physiology parameters at Low and High Light acclimation. (A) Oxygen evolution, 1-qL (fraction of closed reaction centers (RCs)) and Y(NPQ) versus quantum flux for cells acclimated to low light. (B) Oxygen evolution, 1-qL and Y(NPQ) versus quantum flux for cells acclimated to high light. Vertical dashed line indices the quantum flux value where Y(NPQ) exceeds 5% of the F_v_/F_m_ fraction.

**Figure S3.**
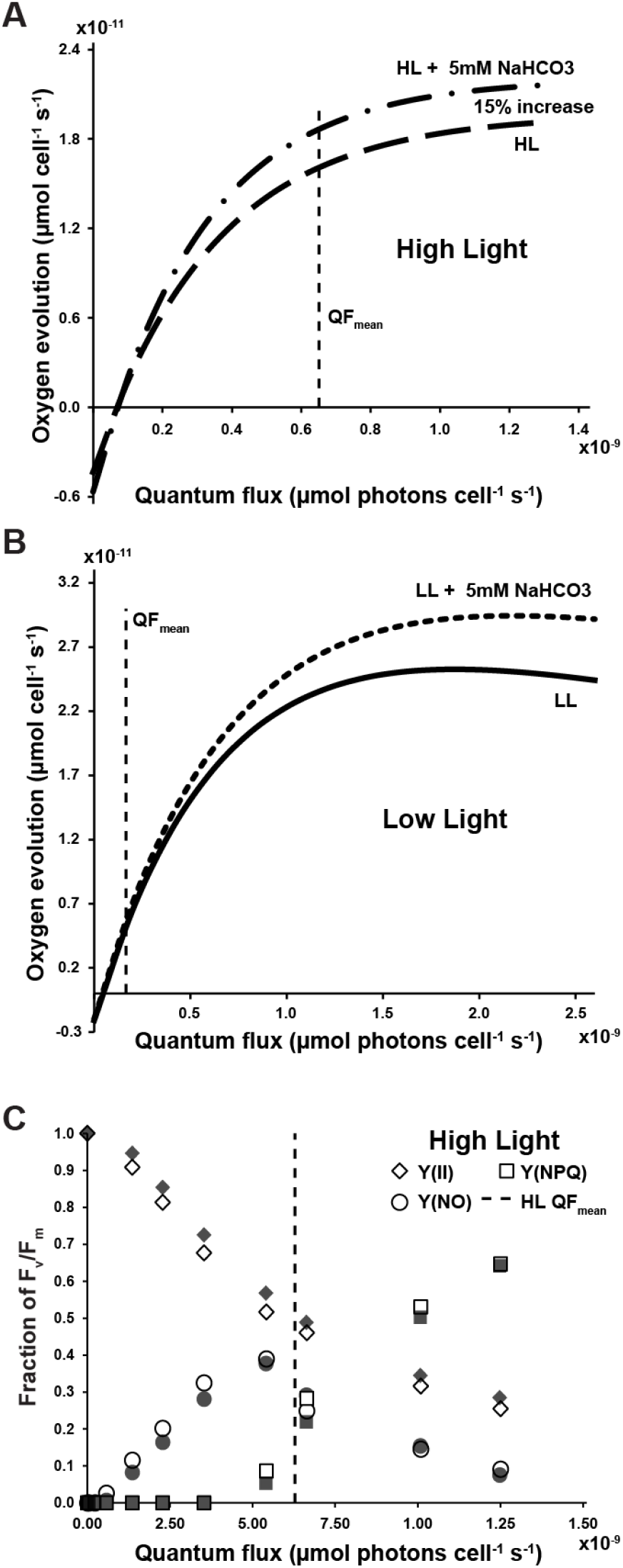
Impact on photosynthetic parameters with and without 5mM bicarbonate. (A) Cell-specific P_O_ versus QF curve for *P. tricornutum* acclimated to high light with and without 5 mM sodium bicarbonate. Vertical dashed lines represent the quantum flux received by the cultures at the experimental irradiance. (B) Cell-specific P_O_ versus QF curve for *P. tricornutum* acclimated to low light with and without 5 mM sodium bicarbonate. Vertical dashed lines represent the quantum flux received by the cultures at the experimental irradiance. (C) Chlorophyll fluorescence measurements (PAM) of *P. tricornutum* acclimated to high light with and without bicarbonate additions. Filled in symbols: +5mM sodium bicarbonate; empty symbols: −5mM sodium bicarbonate. Abbreviations. HL: high light, QF: quantum flux.

**Figure S4.**
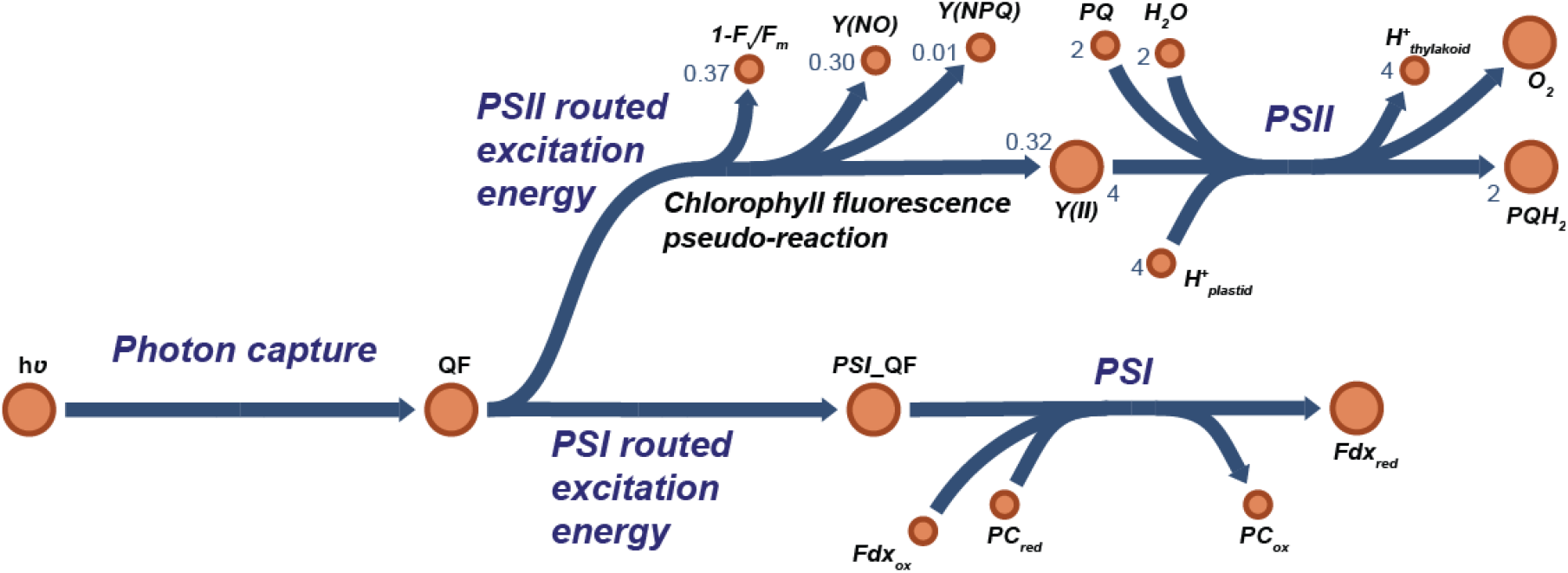
Incorporation of chlorophyll fluorescence parameters in the genome-scale model. Values for the pseudo-reaction are representative of the high light acclimated samples. Abbreviations: hν: photon flux, QF: quantum flux, PSI_QF: quantum flux allocated to photosystem I, Fdxox/red: oxidized/reduced ferredoxin, PCox/red: oxidized/reduced plastocyanin, PQ: oxidized plastoquinone, PQH2: reduced plastoquinone.

**Figure S4.**
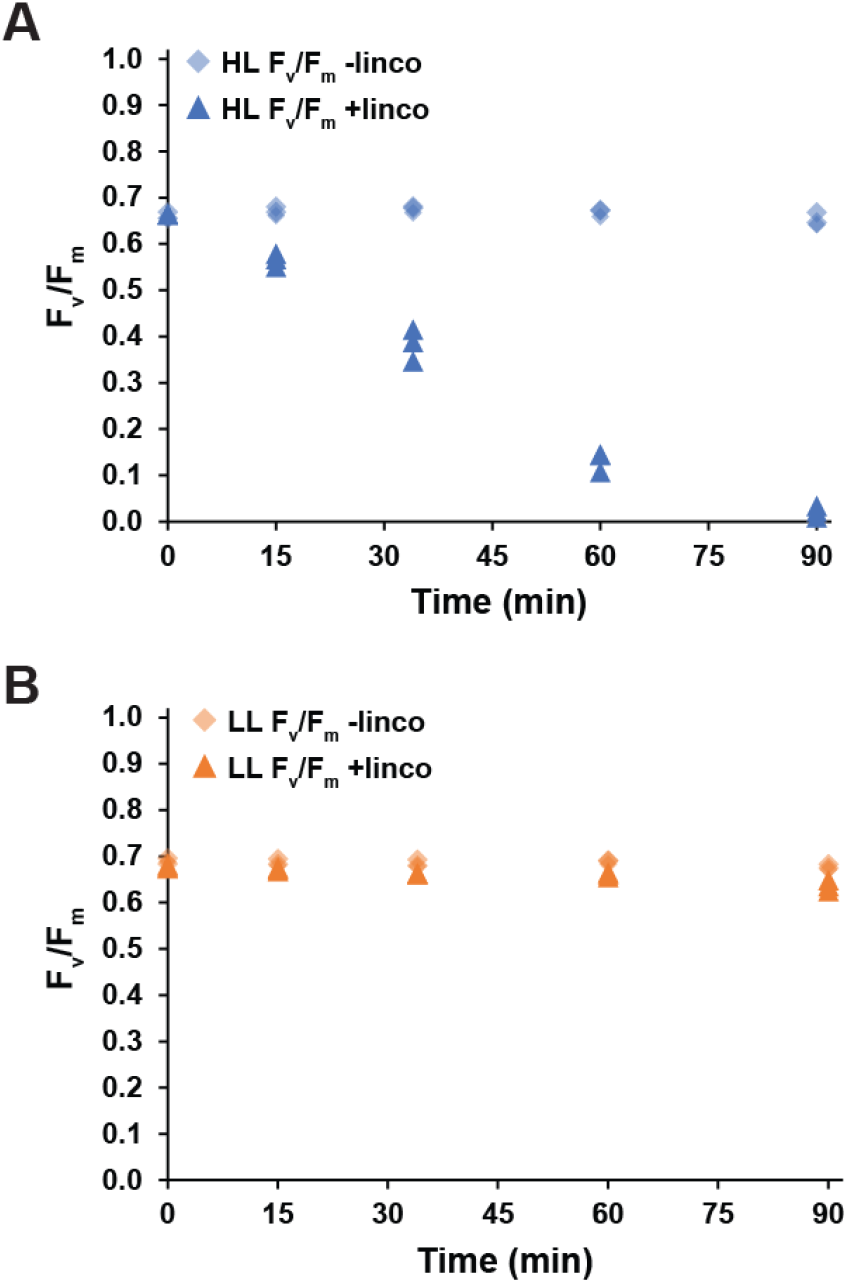
Maximum quantum yield of photosystem II with and without lincomycin treatment. Maximum quantum yield of photosystem II (Fv/Fm), with and without lincomycin treatment, for cells acclimated to (A) high light (n=3) and (B) low light (n=3).

**Figure S5.**
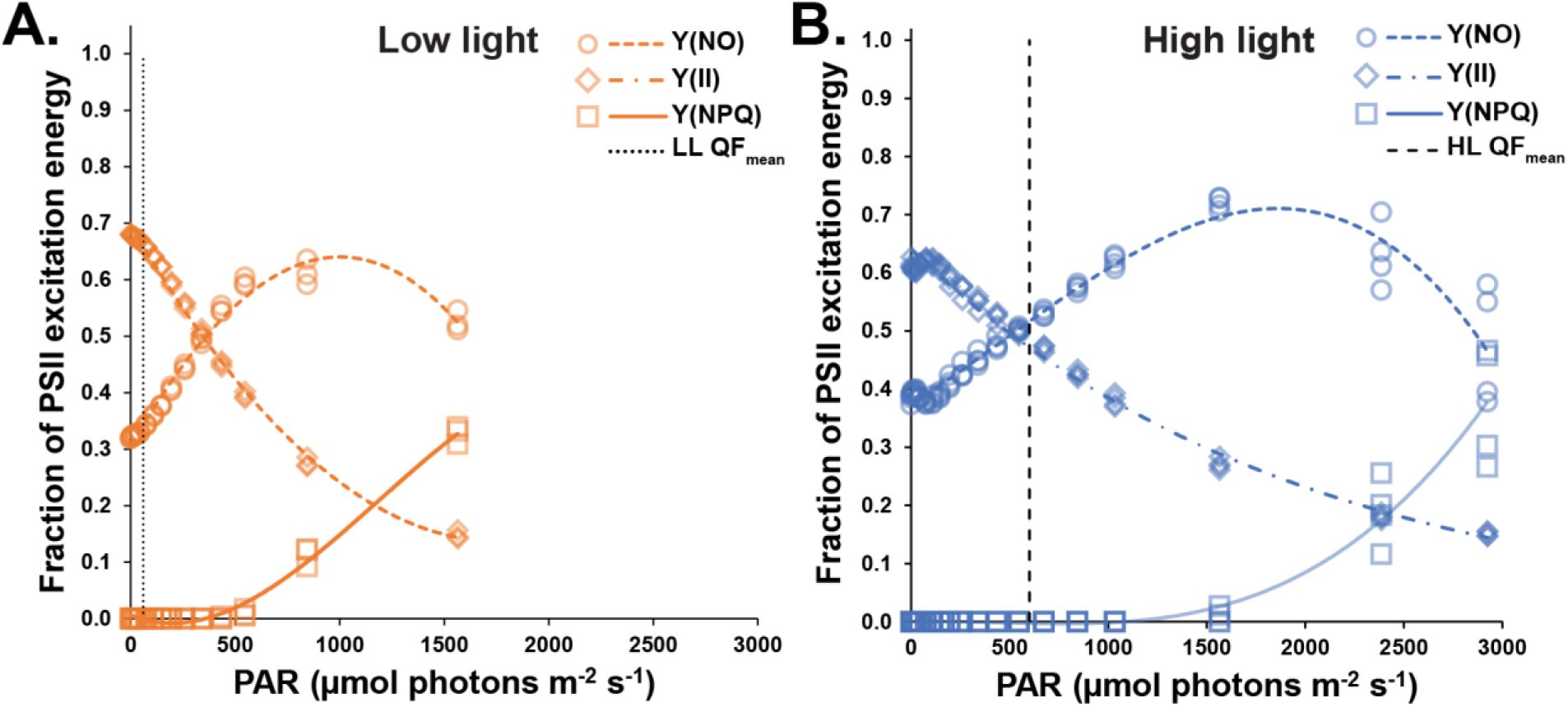
(A) Chlorophyll fluorescence parameters vs. PAR for cells acclimated to low light. (B) Chlorophyll fluorescence parameters vs. PAR for cells acclimated to high light. Vertical dashed lines represent the PAR at the experimental irradiance. Abbreviations and definitions: LL: low light, HL: high light, PAR: Photosynthetically available radiation, Y(II): quantum efficiency of photosystem II, NPQ: non-photochemical quenching, Y(NO): unregulated, non-radiative dissipation of excitation energy. Data based on n=3 biological replicates for LL and n=4 biological replicates for HL.

